# Maintenance DNA methylation is essential for regulatory T cell development and stability of suppressive function

**DOI:** 10.1101/2020.02.03.926949

**Authors:** Kathryn A. Helmin, Luisa Morales-Nebreda, Manuel A. Torres Acosta, Kishore R. Anekalla, Shang-Yang Chen, Hiam Abdala-Valencia, Yuliya Politanska, Paul Cheresh, Mahzad Akbarpour, Elizabeth M. Steinert, Samuel E. Weinberg, Benjamin D. Singer

**Affiliations:** Division of Pulmonary and Critical Care Medicine, Department of Medicine Northwestern University Feinberg School of Medicine Chicago, IL 60611 USA; Department of Surgery Northwestern University Feinberg School of Medicine Chicago, IL 60611 USA; Department of Pathology Northwestern University Feinberg School of Medicine Chicago, IL 60611 USA; Department of Biochemistry and Molecular Genetics Northwestern University Feinberg School of Medicine Chicago, IL 60611 USA; Simpson Querrey Center for Epigenetics Northwestern University Feinberg School of Medicine Chicago, IL 60611 USA

## Abstract

Regulatory T (Treg) cells require Foxp3 expression and induction of a specific DNA hypomethylation signature during development, after which Treg cells persist as a self-renewing population that regulates immune system activation. Whether maintenance DNA methylation is required for Treg cell lineage development and stability and how methylation patterns are maintained during lineage self-renewal remain unclear. Here, we demonstrate that the epigenetic regulator Uhrf1 is essential for maintenance of methyl-DNA marks that stabilize Treg cellular identity by repressing effector T cell transcriptional programs. Constitutive and induced deficiency of Uhrf1 within Foxp3^+^ cells resulted in global yet non-uniform loss of DNA methylation, derepression of inflammatory transcriptional programs, destabilization of the Treg cell lineage, spontaneous inflammation, and enhanced tumor immunity. These findings support a paradigm in which maintenance DNA methylation is required in distinct regions of the Treg cell genome for both lineage establishment and stability of identity and suppressive function.

## Introduction

CD4^+^Foxp3^+^ regulatory T (Treg) cells prevent catastrophic inflammation and facilitate tumor growth by suppressing immune system activation and promoting self-tolerance (1–3). The Foxp3 transcription factor serves as the Treg cell lineage-specifying marker, and Treg cells require constitutive Foxp3 expression to maintain their identity and suppressive function. Mice lacking Treg cells owing to a mutation in the *Foxp3* gene exhibit the scurfy phenotype, succumbing to multi-organ lymphoproliferative inflammation approximately 4 weeks after birth (4, 5). Humans with *FOXP3* mutations develop similar endocrine and enteral inflammation as part of the immunodysregulation, polyendocrinopathy, enteropathy, X-linked (IPEX) syndrome (6). Treg cell dysfunction contributes to the pathogenesis of numerous autoimmune conditions, including systemic lupus erythematosus (7–11) and systemic sclerosis (12). In contrast, modern cancer immunotherapy blocks Treg cell suppressive function to dis-inhibit effector T cell-mediated killing of malignant cells (13–15). Thus, mechanisms involved with both development and stability of the Treg cell lineage represent targets for therapies aimed at amelioration of autoimmune and malignant diseases (3).

Treg cells develop from CD4-single-positive auto-reactive thymocytes that receive signals via CD28/Lck, the IL-2-CD25-Stat5 axis, and T cell receptor engagement by MHC-self-peptide complexes (16). These events induce Foxp3 expression and independently establish a Treg cell-specific cytosine-phospho-guanine (CpG) hypomethylation pattern at certain genomic loci, including the *Foxp3* locus and other loci whose gene products are important for Treg cell lineage identity and suppressive function (17–19). Consistent with the Treg cell requirement for CpG hypomethylation, pharmacologic inhibition of DNA methyltransferase activity is sufficient to induce Foxp3 expression in conventional CD4^+^ T cells and to potentiate Treg cell suppressive function in multiple models of inflammation (20–23). In contrast, conditional constitutive deletion of the DNA methyltransferase Dnmt1 but not Dnmt3a in Treg cells diminishes their numbers and suppressive function (24). Treg cell-specific Dnmt1 deficiency decreases global methyl-CpG content while maintaining the Treg cell-specific CpG hypomethylation pattern at *Foxp3*. In the periphery, TGF-β induces Foxp3 expression in conventional CD4^+^ T cells, although these cells lack the stabilizing DNA hypomethylation landscape that defines thymus-derived Treg cells (17, 25). A minor population of CD4^+^ T cells can promiscuously and transiently express Foxp3, a phenomenon that has been linked to a population of ex-Foxp3 cells that does not represent epigenetic reprogramming of mature Treg cells (26, 27). Conversely, some thymic emigrants referred to as potential Treg cells may harbor the Treg cell-specific CpG hypomethylation pattern but fail to express Foxp3 (28). In adult mice, lineage-tracing studies determined that self-renewal of mature thymus-derived Treg cells maintains the Treg cell lineage even under inflammatory conditions (29). This self-renewal model presupposes that DNA methylation patterns are passed from parent to daughter cell during the self-renewal process. Nevertheless, underlying mechanisms that regulate the stability of the mature Treg cell pool remain unclear, particularly the role of maintenance DNA methylation in lineage stability.

The epigenetic regulator Uhrf1 (ubiquitin-like with plant homeodomain and RING finger domains 1; also known as Np95 in mice and ICBP90 in humans) serves as a non-redundant adapter protein for Dnmt1 during S phase, recruiting Dnmt1 to hemi-methylated DNA to ensure maintenance of CpG methylation patterns as part of multi-protein gene-repressive complexes (30–33). Uhrf1 also regulates *de novo* DNA methylation via recruitment of Dnmt3a and Dnmt3b to chromatin (34, 35). Global homozygous loss of *Uhrf1* results in embryologic lethality (31), phenocopying homozygous loss of *Dnmt1* (36). Conditional loss of Uhrf1 in T cells using a *Cd4*-Cre driver leads to failure of colonic Treg cell proliferation and maturation in response to commensal bacterial colonization (37). Nevertheless, Uhrf1-deficient naïve T cells generated using the *Cd4*-Cre system are able to suppress experimental colitis due to TGF-β-mediated conversion of these cells into an induced Treg cell state (38). The specific role of Uhrf1-mediated maintenance of DNA methylation in Foxp3^+^ Treg cell development and stability of suppressive function remains unknown. Because of Uhrf1’s non-redundant role in recruiting Dnmt1 to hemi-methylated DNA, we hypothesized that constitutive Treg cell-specific Uhrf1 deficiency would phenocopy Treg cell-specific Dnmt1 deficiency and result in failure of thymic Treg cell development. Additionally, based on pharmacologic studies showing augmentation of Foxp3 expression and Treg cell suppressive function upon inhibition of DNA methyltransferase activity, we initially hypothesized that induction of Uhrf1 deficiency in mature Treg cells would augment their suppressive function. Using conditional and chimeric knockout systems, we determined that Uhrf1-mediated DNA methylation is indeed required for thymic Treg cell development. In surprising contrast with our second hypothesis, we used an inducible conditional Uhrf1 knockout system to discover that maintenance DNA methylation at inflammatory gene loci is essential for stabilizing the identity and suppressive function of mature Treg cells.

## Results

### Treg cell-specific deletion of Uhrf1 results in a lethal inflammatory disorder

To test the necessity of maintenance DNA methylation in Treg cell development and function, we generated Treg cell-specific Uhrf1-deficient mice by crossing mice bearing *loxP* sequences at the *Uhrf1* gene locus (*Uhrf1^fl/fl^*) (**Supplemental Figure 1A**) with mice expressing yellow fluorescent protein (YFP) and Cre recombinase driven by the *Foxp3* promoter (*Foxp3^YFP-Cre^*) (39). Crosses of male *Uhrf1^+/fl^Foxp3^YFP-Cre/Y^* mice with female *Uhrf1^fl/fl^Foxp3^+/YFP-Cre^* mice generated F1 pups in statistically Mendelian ratios, although male *Uhrf1^fl/fl^Foxp3^YFP-Cre/Y^* offspring were under-represented at the time of genotyping (approximately 3 weeks of age) (**Supplemental Figure 1B**). *Uhrf1^fl/fl^Foxp3^YFP-Cre^* mice appeared normal at birth but then exhibited spontaneous mortality with a median survival of 28.5 days (**Figure 1A**). Beginning at approximately 3 weeks of age, *Uhrf1^fl/fl^Foxp3^YFP-Cre^* mice were smaller than littermate control mice (*Uhrf1^+/fl^Foxp3^YFP-Cre^*) and displayed scaly skin with cratering and loss of fur (**Supplemental Figure 1C** and **Figure 1B**). Histological examination of the skin revealed infiltration of the dermis and sub-dermis by a large number of inflammatory cells, including lymphocytes and monocytes. Similar to the skin, nearly every internal organ exhibited a mixed cellular infiltrate consisting predominantly of lymphocytes but also monocytes and neutrophils (**Figure 1C**). This striking lymphocytic infiltrate consisted of both CD4^+^ and CD8^+^ T cells (**Figure 1D**), suggesting a lymphocytic inflammatory disorder reminiscent of the scurfy phenotype (4, 5).

**Figure 1.**
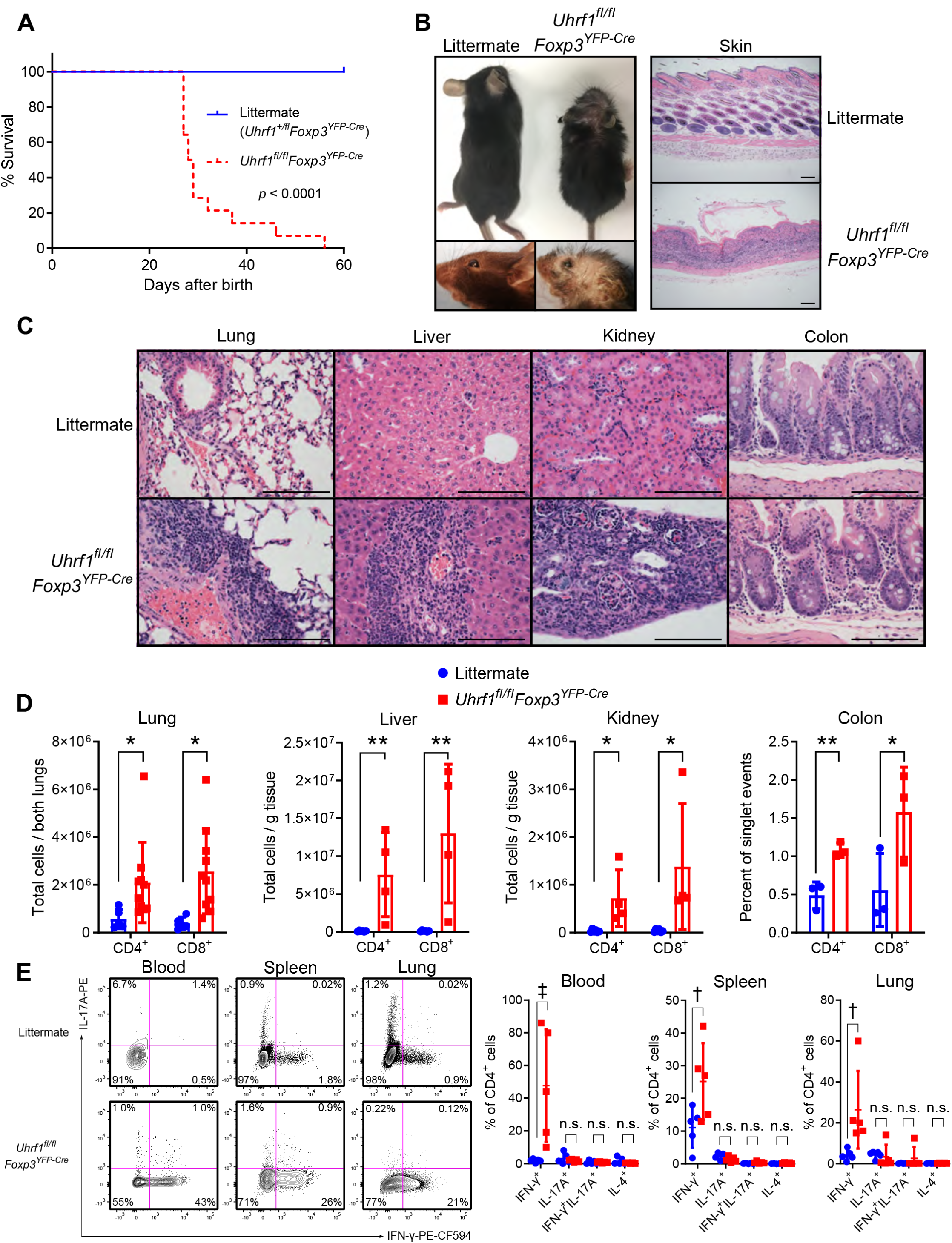
Treg cell-specific Uhrf1-deficient mice spontaneously develop a fatal inflammatory disorder. (**A**) Survival curves of littermate control (*Uhrf1^+/fl^Foxp3^YFP-Cre^*, *n* = 9) and *Uhrf1^fl/fl^Foxp3^YFP-Cre^* (*n* = 14) mice compared using the log-rank (Mantel-Cox) test. (**B**) Gross photographs of 3-4-week-old littermate and *Uhrf1^fl/fl^Foxp3^YFP-Cre^* mice along with photomicrographs of skin. Scale bar represents 100 µm. (**C**) Photomicrographic survey of organ pathology. Scale bar represents 100 µm. (**D**) CD3ε^+^ T cell subsets in selected organs. For lung, *n* = 5 (littermate) and 10 (*Uhrf1^fl/fl^Foxp3^YFP-Cre^*); for liver, *n* = 6 (littermate) and 4 (*Uhrf1^fl/fl^Foxp3^YFP-Cre^*); for kidney, *n* = 6 (littermate) and 4 (*Uhrf1^fl/fl^Foxp3^YFP-Cre^*); for colon, *n* = 3 (littermate and *Uhrf1^fl/fl^Foxp3^YFP-Cre^*). (**E**) Cytokine profile of splenic CD3ε^+^CD4^+^ T cells following *ex vivo* stimulation with phorbol 12-myristate 13-acetate and ionomycin for 4 hours in the presence of brefeldin A. Representative contour plots and summary data are shown. *n* = 5 per group. Summary plots show all data points with mean and standard deviation. * *q* < 0.05, ** *q* < 0.01, † *q* < 0.001, ‡ *q* < 0.0001, n.s. not significant by the two-stage linear step-up procedure of Benjamini, Krieger, and Yekutieli with *Q* = 5%. See Supplemental Table 1 for fluorochrome abbreviations.

**Supplemental Figure 1.**
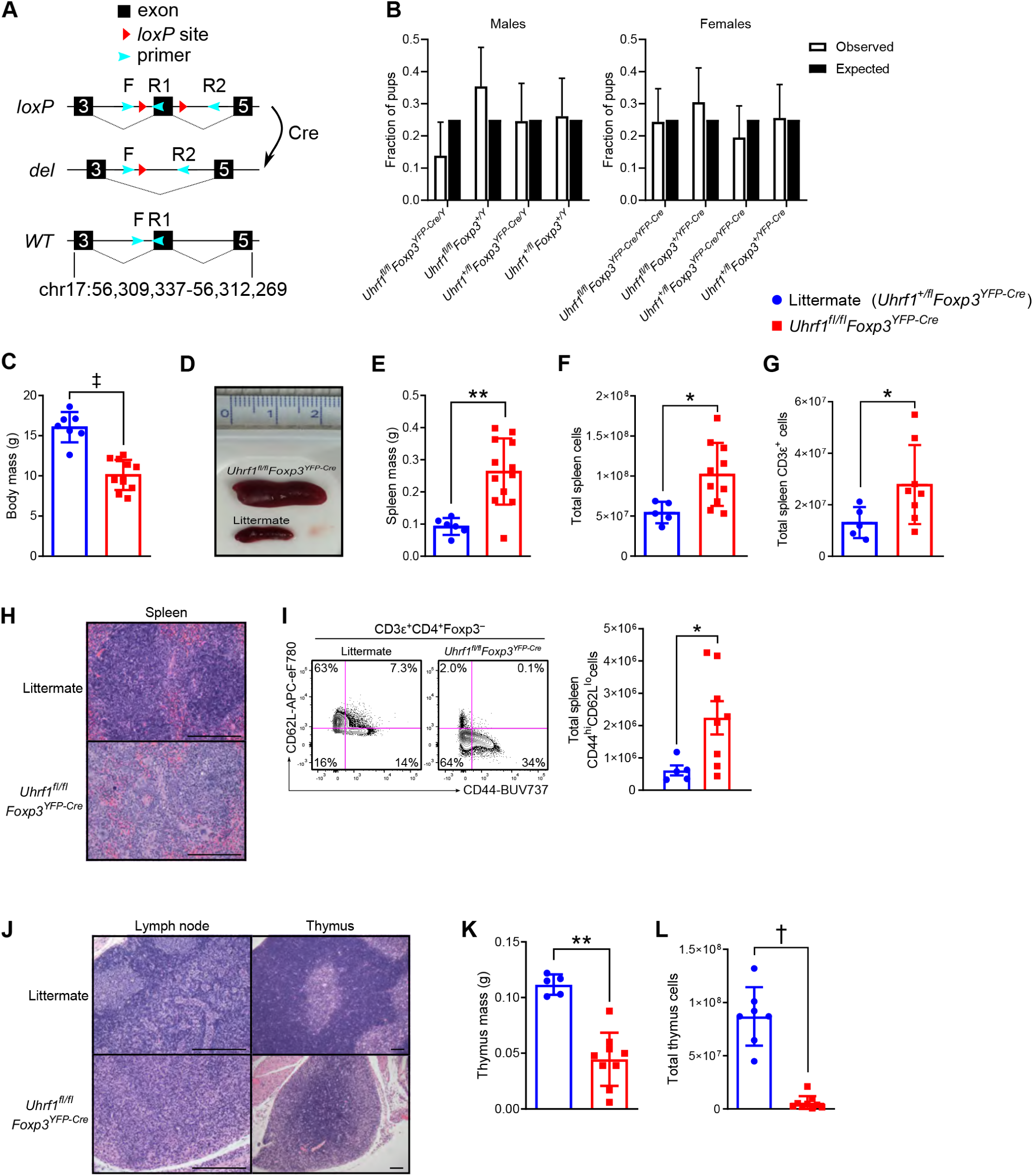
Inflammatory phenotype of 3-4-week-old Treg cell-specific Uhrf1-deficient mice. (**A**) Schematic of the *Uhrf1* locus demonstrating the positions of exons, *loxP* sites, and primers used for genotyping. (**B**) Observed and expected frequencies of F1 pups resulting from crosses of male *Uhrf1^+/fl^Foxp3^YFP-Cre/Y^* mice with female *Uhrf1^fl/fl^Foxp3^+/YFP-Cre^* mice. Chi-square test for goodness of fit *p* = 0.11 for male offspring and 0.57 for female offspring. Chi-square = 6.1 for males and 2.0 for females, both with 3 degrees of freedom. *n* = 65 males and 82 females. (**C**) Body mass of littermate control (*Uhrf1^+/fl^Foxp3^YFP-Cre^*, *n* = 7) and *Uhrf1^fl/fl^Foxp3^YFP-Cre^* (*n* = 11) mice. (**D**) Gross spleen photomicrographs with cm-ruler. (**E**) Splenic masses from littermate (*n* = 6) and *Uhrf1^fl/fl^Foxp3^YFP-Cre^* (*n* = 12) mice. (**F**) Spleen cellularity of littermate (*n* = 5) and *Uhrf1^fl/fl^Foxp3^YFP-Cre^* (*n* = 10) mice. (**G**) Spleen CD3ε^+^ T cell numbers from littermate (*n* = 5) and *Uhrf1^fl/fl^Foxp3^YFP-Cre^* (*n* = 8) mice. (**H**) Photomicrographs of spleen. Scale bar represents 100 µm. (**I**) Activation state of splenic CD3ε^+^CD4^+^Foxp3^−^ cells. Representative contour plots and summary data of CD44^hi^CD62L^lo^ cells are shown. *n* = 5 (littermate) and 8 (*Uhrf1^fl/fl^Foxp3^YFP-Cre^*). (**J**) Photomicrographs of lymph node and thymus. Scale bar represents 100 µm. (**K**) Thymic mass of littermate (*n* = 5) and *Uhrf1^fl/fl^Foxp3^YFP-Cre^* (*n* = 9) mice. (**L**) Thymus cellularity of littermate (*n* = 7) and *Uhrf1^fl/fl^Foxp3^YFP-Cre^* (*n* = 9) mice. Summary plots show all data points with mean and standard deviation except for (B), which shows mean and 95% confidence interval (Wilson/Brown method). * *p* < 0.05, ** *p* < 0.01, † *p* < 0.001, ‡ *p* < 0.0001 by Mann-Whitney test. See Supplemental Table 1 for fluorochrome abbreviations.

Treg cell-specific Uhrf1-deficient mice showed other signs of lymphocyte-driven immune system activation, including splenomegaly and splenic structural disarray characterized by architectural disruption and lymphoid hyperplasia (**Supplemental Figure 1, D-H**). CD3ε^+^CD4^+^ T cells in the spleen displayed an activated profile, exhibiting an increased frequency and total number of CD44^hi^CD62L^lo^ effector T cells (**Supplemental Figure 1I**). Other secondary lymphoid organs also exhibited evidence of immune system activation, including replacement of lymph node germinal centers with a mixed cellular infiltrate and lymphocyte depletion of the thymus, which was also atrophic (**Supplemental Figure 1, J-L**). To better characterize this severe inflammation, we performed intracellular cytokine profiling of CD3ε^+^CD4^+^ T cells from the blood, spleen, and lung. These measurements revealed skewing toward a T helper type 1 (Th1) profile in *Uhrf1^fl/fl^Foxp3^YFP-Cre^* mice, exemplified by a significant increase in the frequency of cells producing interferon-γ (**Figure 1E**). No bacteria were identified in the blood of 3-4-week-old mice by routine culture on LB agar. Collectively, Treg cell-specific deletion of Uhrf1 resulted in severe Th1-skewed inflammation, lymphocytic infiltration of essential organs, thymic atrophy, and mortality at approximately 3-4 weeks of age, suggesting insufficiency or dysfunction of Treg cells as a result of Uhrf1 deficiency.

### Uhrf1 deficiency results in failure of Treg cells to persist after Foxp3 induction in the thymus

We next examined the CD3ε^+^CD4^+^Foxp3^+^ T cell compartment of *Uhrf1^fl/fl^Foxp3^YFP-Cre^* mice to evaluate the effect of Uhrf1 deficiency on Treg cells. Cre-mediated excision effectively deleted *Uhrf1* and revealed the expected pattern of *Uhrf1* genotypes within the sorted *Foxp3*-YFP^−^ and *Foxp3*-YFP^+^ populations of CD3ε^+^CD4^+^ T cells from littermate control mice (*Uhrf1^+/fl^Foxp3^YFP-Cre^*), confirming the Foxp3^+^ cell-exclusive nature of the *Foxp3^YFP-Cre^* construct when paired with a *Uhrf1^fl^* allele (**Supplemental Figure 2A**). At 3-4 weeks of age, *Uhrf1^fl/fl^Foxp3^YFP-Cre^* mice exhibited a profound deficiency of Treg cells, with Foxp3^+^ cells constituting < 1% of CD3ε^+^CD4^+^ T cells in the blood, spleen, and lung (**Figure 2A**). Examination of Foxp3^+^ Treg cells marked by high expression of CD25 (IL-2Rα) as a fraction of CD3ε^+^CD4^+^ T cells further highlighted the Treg cell deficiency in *Uhrf1^fl/fl^Foxp3^YFP-Cre^* mice (**Figure 2B**). Total numbers of Foxp3^+^ Treg cells were likewise diminished in the spleen and lung, representative of secondary lymphoid and parenchymal organs, respectively (**Figure 2C**).

**Supplemental Figure 2.**
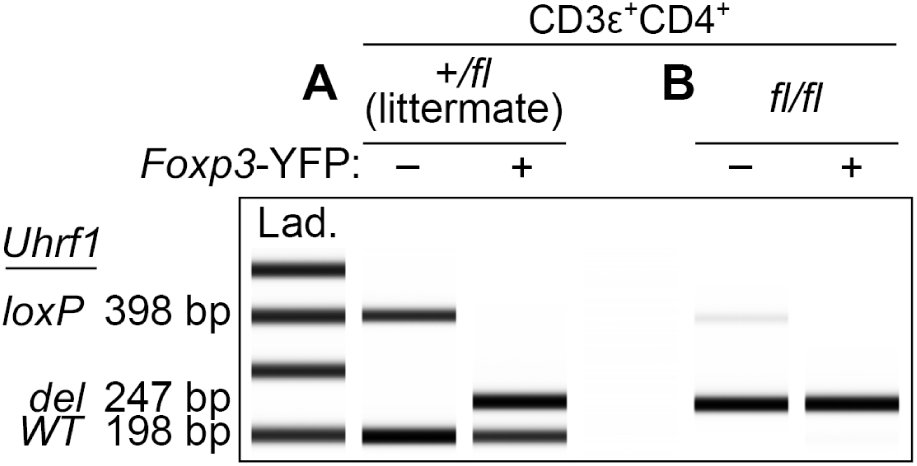
Genotype of sorted splenic CD3ε^+^CD4^+^ T cell populations. (**A and B**) Agilent TapeStation image of PCR products generated using the displayed primer scheme shown in Supplemental Figure 1A. Splenic CD3ε^+^CD4^+^ cells were sorted based on *Foxp3*-YFP status from 3-4-week-old *Uhrf1^+/fl^Foxp3^YFP-Cre^* littermate control mice (A) and *Uhrf1^fl/fl^Foxp3^YFP-Cre^* mice (B). Image is representative of three independent experiments. Lad., ladder.

**Figure 2.**
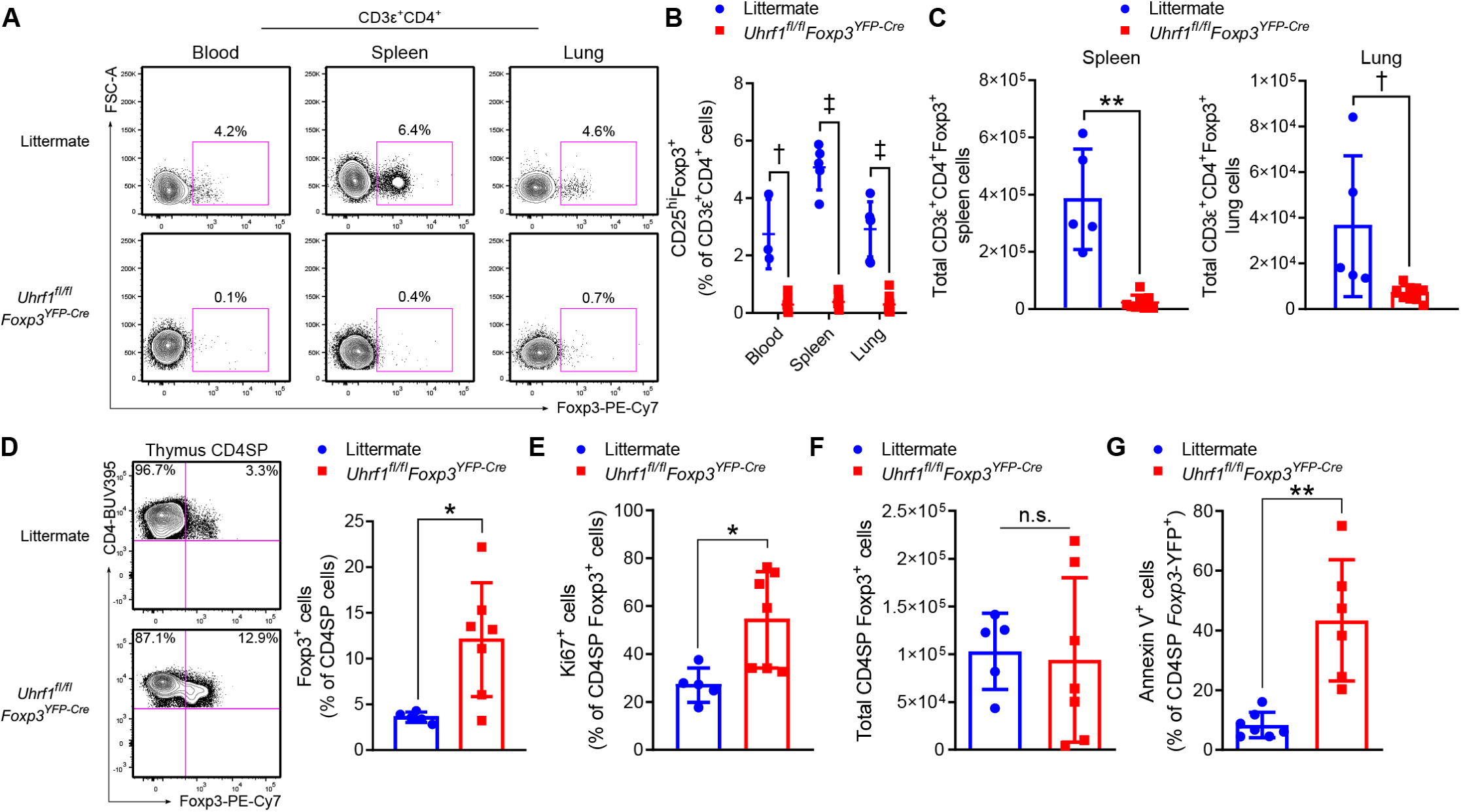
Treg cells are reduced in the periphery of 3-4-week-old Treg cell-specific Uhrf1-deficient mice. (**A**) Foxp3^+^ cells as a frequency of CD3ε^+^CD4^+^ cells in blood, spleen, and lung shown as representative flow cytometry contour plots. (**B**) CD25^hi^Foxp3^+^ cells expressed as a percentage of CD3ε^+^CD4^+^ cells for blood, spleen, and lung. For blood, *n* = 3 (*Uhrf1^+/fl^Foxp3^YFP-Cre^* littermate) and 7 (*Uhrf1^fl/fl^Foxp3^YFP-Cre^*); for spleen, *n* = 5 (littermate) and 9 (*Uhrf1^fl/fl^Foxp3^YFP-Cre^*); for lung, *n* = 6 (littermate) and 10 (*Uhrf1^fl/fl^Foxp3^YFP-Cre^*). (**C**) Total spleen and lung CD3ε^+^CD4^+^Foxp3^+^ cells. For spleen, *n* = 5 (littermate) and 8 (*Uhrf1^fl/fl^Foxp3^YFP-Cre^*); for lung, *n* = 5 (littermate) and 10 (*Uhrf1^fl/fl^Foxp3^YFP-Cre^*). (**D**) Thymic Foxp3^+^ cell frequency. Representative flow cytometry contour plots of CD4-single-positive (SP) thymocytes from littermate and *Uhrf1^fl/fl^Foxp3^YFP-Cre^* mice. Foxp3^+^ cells are shown as a percentage of the CD4SP population. *n* = 5 (littermate) and 7 (*Uhrf1^fl/fl^Foxp3^YFP-Cre^*). (**E**) Proliferating (Ki67^+^) cells as a percentage of CD4SP Foxp3^+^ thymocytes. *n* = 5 (littermate) and 7 (*Uhrf1^fl/fl^Foxp3^YFP-Cre^*). (**F**) Total thymic Treg cells. *n* = 5 (littermate) and 7 (*Uhrf1^fl/fl^Foxp3^YFP-Cre^*). (**G**) Apoptotic (annexin V^+^) cells as a percentage of CD4SP Foxp3^+^ thymocytes. *n* = 7 (littermate) and 6 (*Uhrf1^fl/fl^Foxp3^YFP-Cre^*). Summary plots show all data points with mean and standard deviation. * *p* < 0.05, ** *p* < 0.01, † *q* or *p* < 0.001, ‡ *q* < 0.0001, n.s. not significant by the two-stage linear step-up procedure of Benjamini, Krieger, and Yekutieli with *Q* = 5% (B) or Mann-Whitney test (C-G). FSC-A, forward scatter area; see Supplemental Table 1 for fluorochrome abbreviations.

The *Foxp3^YFP-Cre^* construct is first expressed at sustained levels at the thymic Treg cell stage of development (19). Accordingly, we examined the fraction of Foxp3^+^ cells among CD4-single-positive cells in the thymus and found that thymic Treg cells were increased in frequency among *Uhrf1^fl/fl^Foxp3^YFP-Cre^* mice (**Figure 2D**) and displayed increased proliferation (**Figure 2E**). Nevertheless, total thymic Treg cell numbers were similar between littermates and *Uhrf1^fl/fl^Foxp3^YFP-Cre^* mice (**Figure 2F**), likely due of the severe thymic atrophy observed among *Uhrf1^fl/fl^Foxp3^YFP-Cre^* animals (see Supplemental Figure 1, J-L). This observation excludes the possibility of thymic accumulation or trapping of Treg cells as a cause of their peripheral deficiency. To examine the possibility that loss of Uhrf1 at the time of *Foxp3* expression triggers apoptosis, we measured annexin V staining on thymic Treg cells and found an increased frequency of annexin V^+^ thymic Treg cells in *Uhrf1^fl/fl^Foxp3^YFP-Cre^* mice compared with littermate control mice (**Figure 2G**). We then examined the genotype of the sorted *Foxp3*-YFP^−^ and *Foxp3*-YFP^+^ populations from *Uhrf1^fl/fl^Foxp3^YFP-Cre^* mice. The *Foxp3*-YFP^+^ population contained a *Uhrf1-deleted* sequence, confirming Cre-mediated deletion of *Uhrf1* and excluding the possibility that the small population of *Foxp3*-YFP^+^ cells escaped Cre-mediated recombination (**Supplemental Figure 2B**). To our surprise, however, the *Foxp3*-YFP^−^ population contained a minor *Uhrf1-loxP* signal and a dominant *Uhrf1-deleted* sequence, suggesting a population of ex-Foxp3 cells that once expressed the *Foxp3^YFP-Cre^* construct but lost *Foxp3* expression (and thus YFP^+^ status) shortly after the thymic Foxp3^+^ Treg cell stage of development. In summary, loss of Uhrf1 at the thymic Treg cell stage of development resulted in failure of Treg cells to develop into a robust population, indicating the necessity of Uhrf1 in developmental stabilization of the Treg cell lineage.

### Uhrf1 deficiency leads to a cell-autonomous and inflammation-independent decrease in Treg cells without altering Foxp3 expression

To establish that the lack of Treg cells in Treg cell-specific Uhrf1-deficient animals is a cell-autonomous effect of Uhrf1 deficiency and not due to the profound inflammation observed in *Uhrf1^fl/fl^Foxp3^YFP-Cre^* mice, we generated Uhrf1 chimeric knockout animals—female mice that are homozygous for the *Uhrf1^fl^* allele and heterozygous for the X-linked *Foxp3^YFP-Cre^* allele. Following random inactivation of the X chromosome in these mice, the Treg cell compartment contains a mix of Uhrf1-sufficient *Foxp3*-YFP^−^ and Uhrf1-deficient *Foxp3*-YFP^+^ cells. Uhrf1 chimeric knockout mice (*Uhrf1^fl/fl^Foxp3^+/YFP-Cre^*) displayed a significant decrease in *Foxp3*-YFP^+^ cells compared with control mice (*Uhrf1^+/+^Foxp3^YFP-Cre/YFP-Cre^*) (**Figure 3A**). Consistent with the chimeric construct, CD25 expression was distributed throughout the *Foxp3*-YFP^−^ and small *Foxp3*-YFP^+^ compartments of Uhrf1 chimeric knockout animals but co-segregated with *Foxp3*-YFP expression in control mice (**Figure 3B**). Despite the dearth of *Foxp3*-YFP^+^ cells in Uhrf1 chimeric knockout animals, average per-cell Foxp3 protein expression was similar to control mice (**Supplemental Figure 3A**). Uhrf1 chimeric knockout mice appeared grossly normal, did not experience overt spontaneous inflammation, and did not display increased spleen mass, cellularity, or frequency of CD44^hi^CD62L^lo^ effector T cells (**Supplemental Figure 3, B-D**). Taken together, these results demonstrate that Treg cell-specific Uhrf1 deficiency causes an inflammation-independent lack of Treg cells in a cell-autonomous fashion. These data also suggest that Uhrf1-suficient cells are able to fill the Treg cell niche in the presence of a population of Uhrf1-deficient cells.

**Figure 3.**
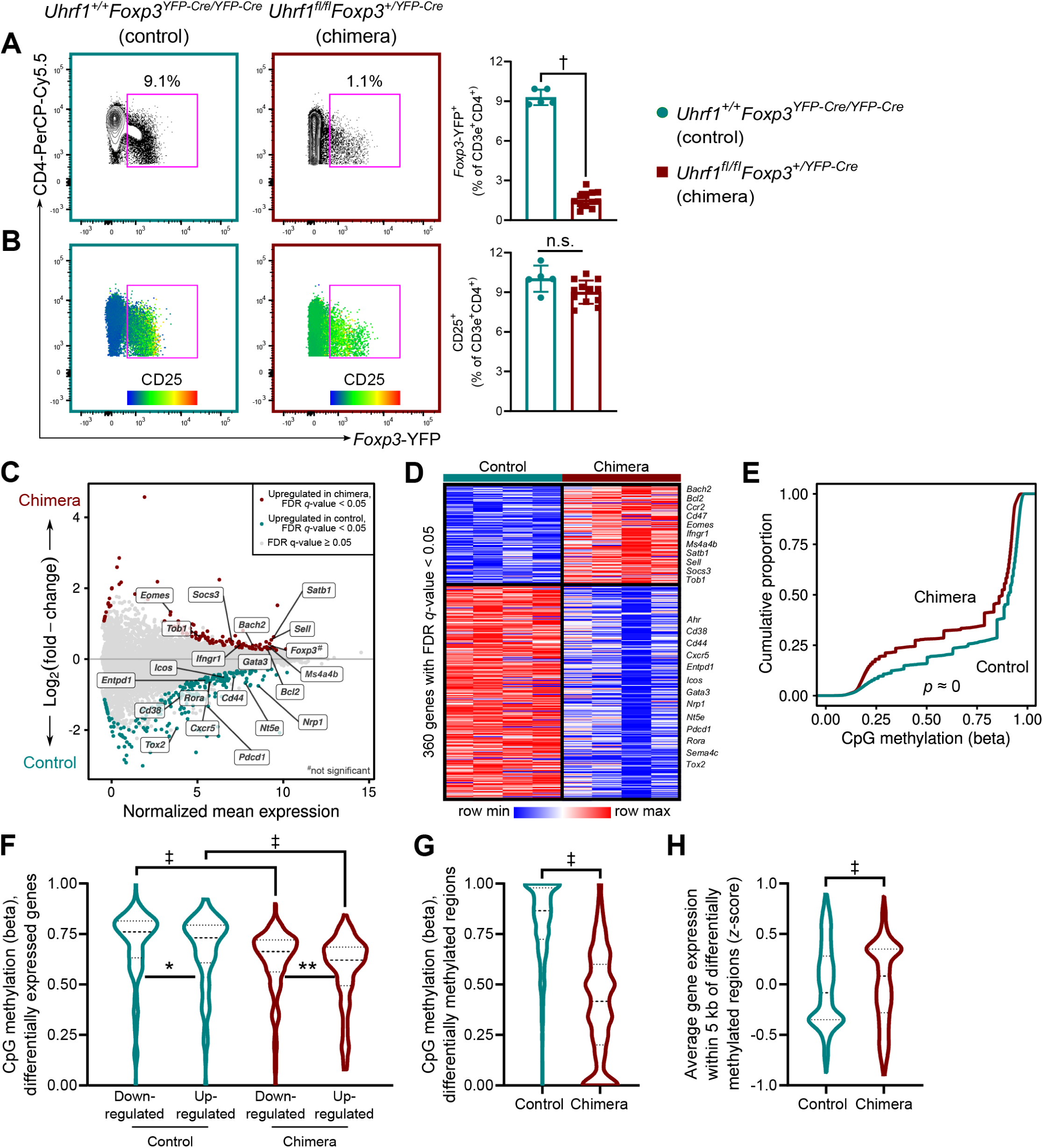
Treg cell-specific Uhrf1 chimeric knockout mice reveal the Treg cell-autonomous and inflammation-independent effects of Uhrf1 deficiency. (**A**) Representative contour plots and quantification showing *Foxp3*-YFP^+^ cells as a frequency of splenic CD3ε^+^CD4^+^ cells from 8-week-old female *Uhrf1^+/+^Foxp3^YFP-Cre/YFP-Cre^* (control) and *Uhrf1^fl/fl^Foxp3^+/YFP-Cre^* (chimeric) mice. (**B**) CD25 heat map overlaid on populations shown in (A); the associated graph quantifies CD25^+^ cells as a percentage of splenic CD3ε^+^CD4^+^ cells. (**C**) MA plot comparing gene expression of splenic CD3ε^+^CD4^+^CD25^hi^*Foxp3*-YFP^+^ cells from control mice (Uhrf1-sufficient cells) and Uhrf1 chimeric knockout mice (Uhrf1-deficient cells). Genes of interest are annotated. (**D**) *K*-means clustering of 360 genes with an FDR *q*-value < 0.05 comparing the cell populations from (C) with *k* = 2 and scaled as *z*-score across rows. Genes of interest are annotated. (**E**) Cumulative distribution function plot of 1.4 x 10^6^ well-observed CpGs. CpG methylation is expressed as beta scores with 0 representing unmethylated and 1 representing fully methylated; a shift in the cumulative distribution function up and to the left represents relative hypomethylation. (**F**) CpG methylation at the loci of differentially expressed genes (defined as the gene body ± 2 kb). (**G**) CpG methylation at 1,335 differentially methylated regions. (**H**) Average gene expression (*z*-score) at 567 gene loci within 5 kb of differentially methylated regions. *n* = 5 (control) and 12 (chimera) for (A) and (B) and 4 mice per group for (C-F). Summary plots show all data points with mean and standard deviation; violin plots show median and quartiles. * *q* < 0.05, ** *q* < 0.01, † *p* < 0.001, ‡ *q* or *p* < 0.0001, n.s. not significant by Mann-Whitney test (A and B), a mixed-effects analysis with the two-stage linear step-up procedure of Benjamini, Krieger, and Yekutieli with *Q* = 5% (F), or Kolmogorov-Smirnov test for cumulative distributions (G and H). The approximate *p*-value resulting from a Kolmogorov-Smirnov test for cumulative distributions is shown in (E). See Supplemental Table 1 for fluorochrome abbreviations.

**Supplemental Figure 3.**
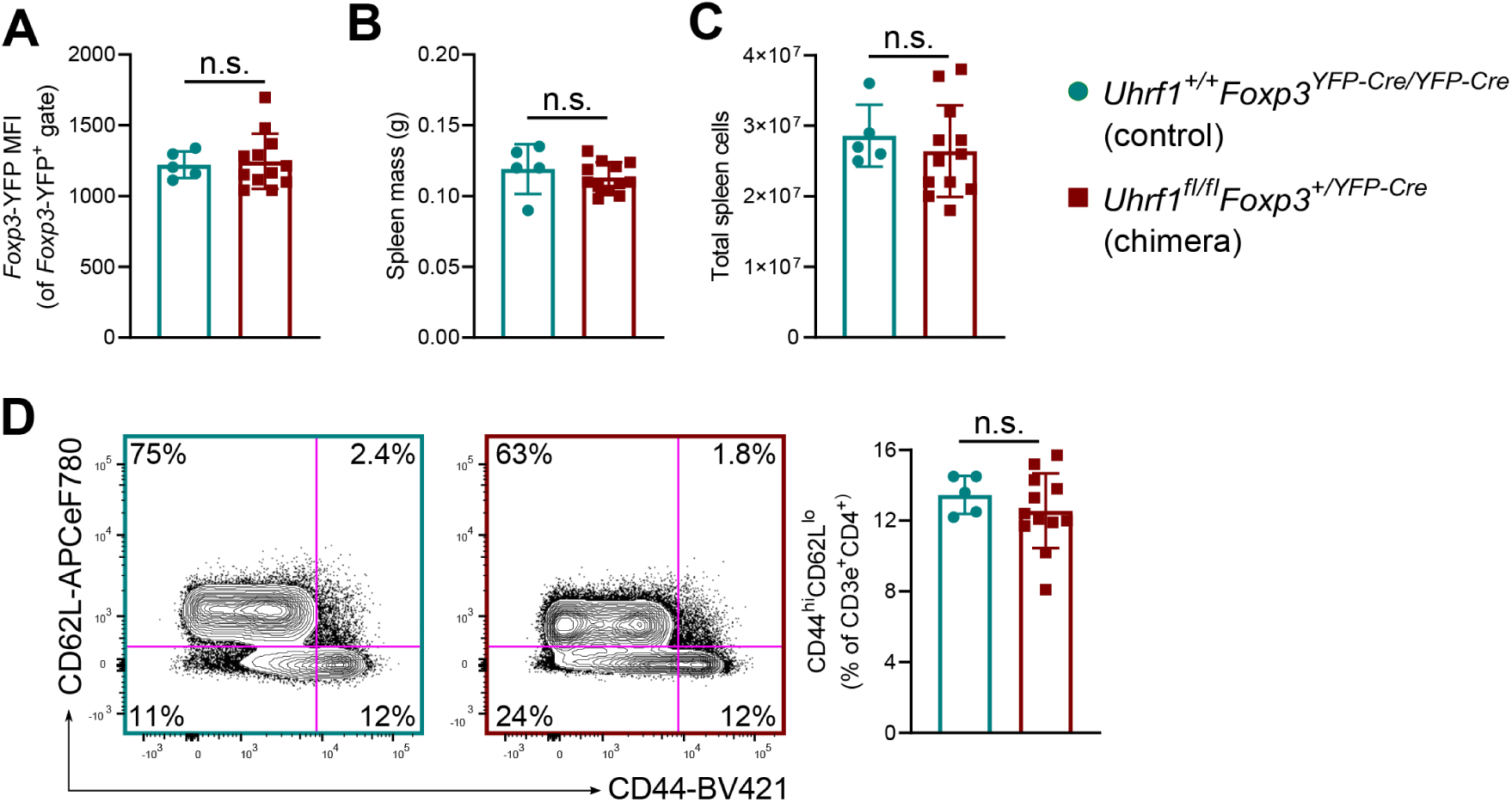
Extended phenotype of Treg cell-specific Uhrf1 chimeric knockout mice. (**A**) *Foxp3*-YFP mean fluorescence intensity (MFI) within the splenic *Foxp3*-YFP^+^ population of 8-week-old female *Uhrf1^+/+^Foxp3^YFP-Cre/YFP-Cre^* (control) and *Uhrf1^fl/fl^Foxp3^+/YFP-Cre^* (chimeric) mice. (**B**) Splenic mass of control and chimeric mice. (**C**) Spleen cellularity of control and chimeric mice. (**D**) Activation state of splenic CD3ε^+^CD4^+^ cells. Representative contour plots and summary data of CD44^hi^CD62L^lo^ cells are shown. Summary plots show all data points with mean and standard deviation*. n* = 5 (control) and 12 (chimera). n.s. not significant by Mann-Whitney test. See Supplemental Table 1 for fluorochrome abbreviations.

### Foxp3^+^ cells from Uhrf1 chimeric knockout mice exhibit downregulation of genes associated with Treg cell suppressive function and loss of DNA methylation while preserving Foxp3 locus expression and methylation patterning

We took advantage of the Uhrf1 chimeric knockout system to investigate inflammation-independent mechanisms of impaired suppressive function in Uhrf1-deficient Treg cells. Accordingly, we performed transcriptional profiling of CD3ε^+^CD4^+^CD25^hi^*Foxp3*-YFP^+^ cells isolated from control and Uhrf1 chimeric knockout mice. RNA-sequencing followed by an unsupervised analysis revealed differential expression of 360 genes with a false-discovery rate (FDR) *q*-value < 0.05 (**Figure** 3, C **and** D). Genes that were downregulated in cells from chimeric knockout animals included those classically associated with Treg cell activation and suppressive function: *Ahr*, *Cd38*, *Cd44*, *Entpd1*, *Icos*, *Gata3*, *Nrp1*, *Nt5e*, *Pdcd1*, *Sema4c*, and *Tox2*. Genes upregulated in knockout cells included *Bach2*, *Bcl2*, *Ccr2*, *Cd47*, *Eomes*, *Ifngr1*, *Ms4a4b*, *Satb1*, *Sell*, *Socs3*, and *Tob1*—many of which are associated with impaired Treg suppressive function and gain of effector T cell function. Importantly, *Foxp3* expression was not significantly affected by Uhrf1 deficiency (log2(fold-change) = 0.39, FDR *q*-value = 0.06). Gene Set Enrichment Analysis using a list of genes canonically associated with Treg cell identity (40) revealed significant negative enrichment within cells from chimeric knockout mice (**Supplemental Figure 4A**). These findings in non-inflamed Uhrf1 chimeric knockout mice confirm that Uhrf1 deficiency results in loss of the canonical Treg cell transcriptional program without significantly affecting levels of *Foxp3* expression in Foxp3^+^ cells.

**Supplemental Figure 4.**
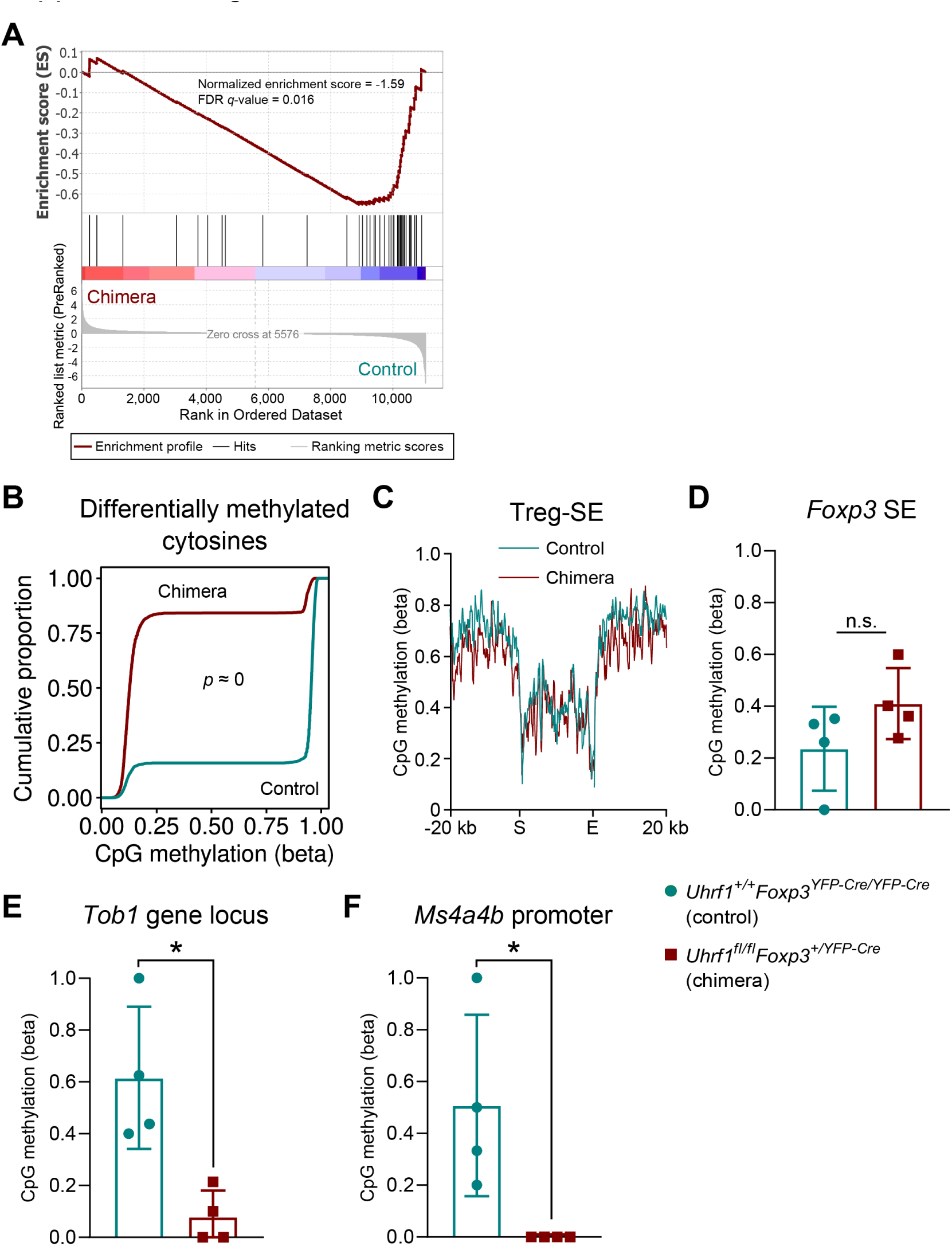
Extended molecular analysis of Treg cell-specific Uhrf1 chimeric knockout mice. (**A**) Gene set enrichment plot testing for enrichment of a Treg cell-defining gene set (40) with genes ordered by log2(fold-change) in average expression comparing CD3ε^+^CD4^+^CD25^hi^*Foxp3*-YFP^+^ cells from Uhrf1 chimeric knockout mice against control mice. The normalized enrichment score (NES) and FDR *q*-value associated with this test are shown. (**B**) Cumulative distribution function plot of 4,708 CpGs with FDR *q*-value < 0.05 compared using a Wald test for beta-binomial distributions. (**C**) Metagene analysis of CpG methylation across the start (S) through the end (E) of Treg cell-specific super-enhancer elements (Treg-SE) as defined in reference (19). (**D-F**) CpG methylation status of the *Foxp3* locus super-enhancer (D, chrX:7,565,077-7,587,439), the *Tob1* gene locus (E, chr11:94,209,454-94,217,495), and the *Ms4a4b* promoter (F, chr19:11,442,553-11,444,553). *n* = 4 mice per group. Summary plots show all data points with mean and standard deviation. The approximate *p*-value resulting from a Kolmogorov-Smirnov test for cumulative distributions is shown in (B). * *p* < 0.05, n.s. not significant by Mann-Whitney test (D-F).

Genome-wide CpG methylation profiling with modified reduced representation bisulfite sequencing (mRRBS) revealed a striking global hypomethylation pattern within CD3ε^+^CD4^+^CD25^hi^*Foxp3*-YFP^+^ cells from Uhrf1 chimeric knockout mice compared with control mice (**Figure 3E** and **Supplemental Figure 4B**). CpG methylation was specifically reduced at the loci of differentially expressed genes in cells from chimeric knockout mice, particularly at upregulated loci (**Figure 3F**). We next used an unsupervised procedure to define 1,335 differentially methylated regions (DMRs); these DMRs were significantly hypomethylated in cells from chimeric knockout mice compared with control mice (**Figure 3G**). Gene expression was increased at loci near these DMRs (**Figure 3H**), consistent with loss of Uhrf1-mediated DNA methylation leading to derepression of these loci. Importantly, CpG methylation across Treg cell-specific super-enhancer elements and the super-enhancer at the *Foxp3* locus, which contains the canonical *Foxp3* conserved non-coding sequences (19), was unaffected by Uhrf1 deficiency (**Supplemental Figure 4, C and D**). Examination of gene loci that displayed hypomethylation and upregulation in knockout cells compared with control cells included genes associated with impaired Treg suppressive function and gain of effector T cell function: *Tob1* (41) and the promoter region of *Ms4a4b* (42) (**Supplemental Figure 4, E and F**). Altogether, deficiency of Uhrf1 in Treg cells resulted in loss of the Treg cell lineage-defining transcriptional signature, disruption of DNA methylation patterning, and derepression of effector programs.

### Uhrf1 is required for stability of Treg cell suppressive function

Thymus-derived Treg cells self-renew to maintain the Treg cell lineage (29). Accordingly, we sought to understand how Treg cell-specific loss of Uhrf1 affects the stability of both the developing and mature Treg cell pools. To that end, we generated inducible, Treg cell-specific, *Foxp3* lineage-traceable, Uhrf1-deficient mice (**Figure 4A**). These mice—which we refer to as i*Uhrf1^fl/fl^* mice—are homozygous for the *Uhrf1^fl^* allele, an inducible *Foxp3*-Cre driver with a GFP label (*Foxp3^eGFP-CreERT2^*), and a *loxP*-flanked stop codon upstream of the red florescent protein tdTomato driven by a CAG promoter at the open *ROSA26* locus (*ROSA26Sor^CAG-tdTomato^*). We fed tamoxifen chow to juvenile 4-week-old i*Uhrf1^fl/fl^* mice, which are still developing their Treg cell pool, as well as adult 8-week-old i*Uhrf1^fl/fl^* mice that have established a mature Treg cell lineage with minimal input from thymic emigrants (29). Following tamoxifen administration, both 4- and 8-week-old i*Uhrf1^fl/fl^* mice lost weight compared with i*Uhrf1^+/+^* control mice (**Supplemental Figure 5A and Figure 4B**), suggesting the onset of inflammation in both juvenile and adult i*Uhrf1^fl/fl^* mice. As further evidence of inflammation, both 4- and 8-week-old i*Uhrf1^fl/fl^* mice developed splenomegaly after 4 weeks of tamoxifen (**Supplemental Figure 5B** and **Figure 4C**). Similar to *Uhrf1^fl/fl^Foxp3^YFP-Cre^* mice, 8-week-old i*Uhrf1^fl/fl^* mice fed tamoxifen for 4 weeks exhibited disrupted splenic architecture with distortion of germinal centers by lymphocyte expansion and infiltration with other inflammatory cells (**Figure 4D**). A histological survey of the skin and internal organs of these i*Uhrf1^fl/fl^* mice revealed a phenotype reminiscent of the spontaneous lymphocytic inflammatory process observed in *Uhrf1^fl/fl^Foxp3^YFP-Cre^* mice (**Figure 4E**), albeit less severe, as these i*Uhrf1^fl/fl^* mice were not approaching the moribund status that *Uhrf1^fl/fl^Foxp3^YFP-Cre^* mice displayed at 3-4 weeks of age.

**Figure 4.**
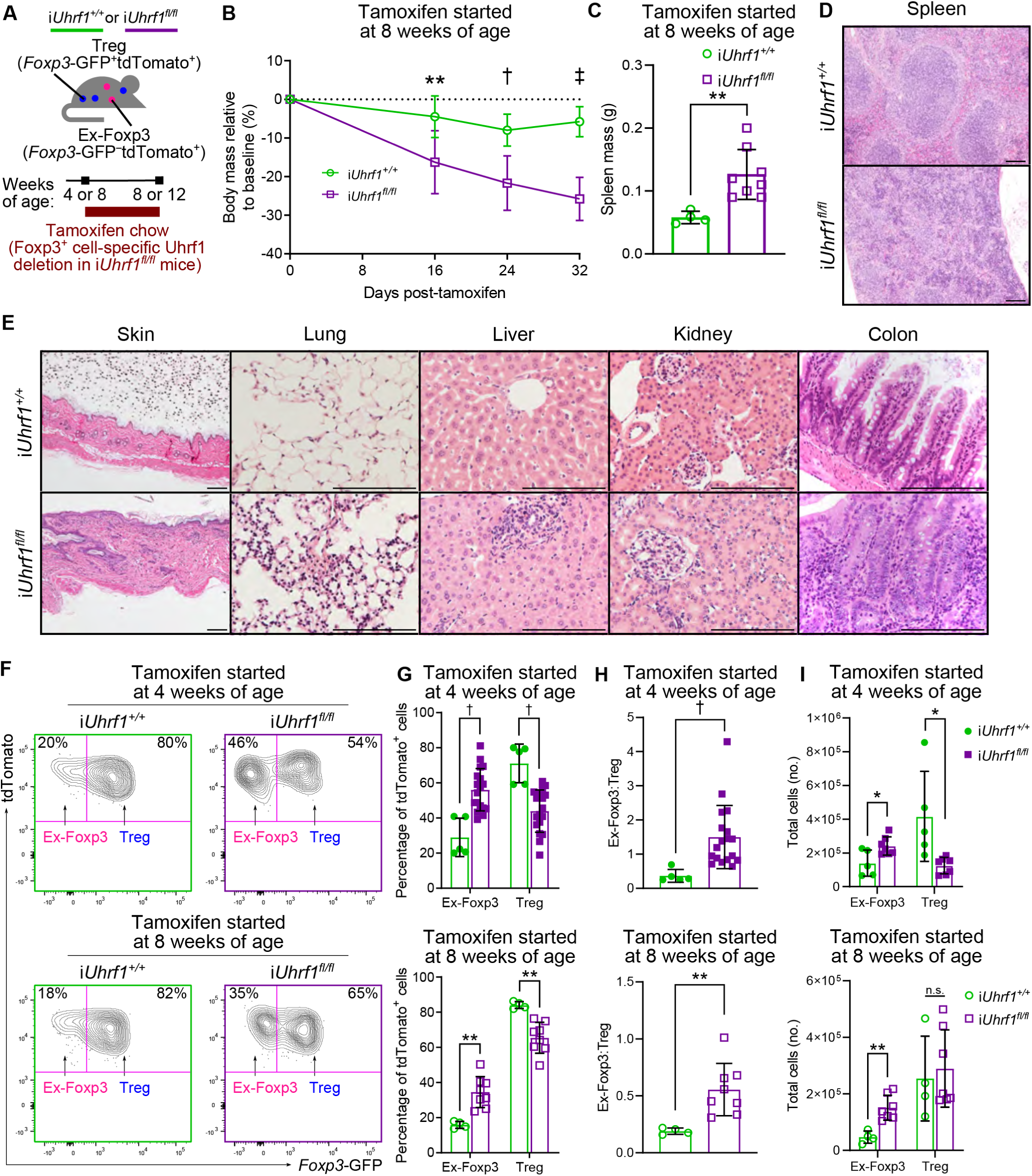
Induced loss of Uhrf1 in Treg cells results in spontaneous inflammation and generation of ex-Foxp3 cells. (**A**) Schematic of experimental design. (**B**) Body mass curves for i*Uhrf1^+/+^* and i*Uhrf1^fl/fl^* mice initiated on tamoxifen chow at 8 weeks of age. *n* = 4 (i*Uhrf1^+/+^*) and 14 (i*Uhrf1^fl/fl^*). (**C**) Spleen masses for i*Uhrf1^+/+^* and i*Uhrf1^fl/fl^* mice initiated on tamoxifen at 8 weeks of age. *n* = 4 (i*Uhrf1^+/+^*) and 8 (i*Uhrf1^fl/fl^*). (**D**) Photomicrographs of spleen pathology from 8-week-old mice after 4 weeks of tamoxifen. Scale bar represents 100 µm. (**E**) Photomicrographic survey of organ pathology from 8-week-old mice after 4 weeks of tamoxifen. Scale bar represents 100 µm. (**F**) Representative flow cytometry contour plots gated on splenic CD3ε^+^CD4^+^tdTomato^+^ cells showing the percentage of ex-Foxp3 (*Foxp3*-GFP^−^) and Treg (*Foxp3*-GFP^+^) cells after 4 weeks of tamoxifen started at either 4 or 8 weeks of age. (**G**) Summary data of the percentage of ex-Foxp3 and Treg cells within the tdTomato^+^ population for the experiment shown in (F). (**H**) Ratio of ex-Foxp3 to Treg cells after 4 weeks of tamoxifen started at either 4 or 8 weeks of age. For mice started on tamoxifen at 4 weeks of age, *n* = 5 (i*Uhrf1^+/+^*) and 18 (i*Uhrf1^fl/fl^*); for mice started on tamoxifen at 8 weeks of age, *n* = 4 (i*Uhrf1^+/+^*) and 8 (i*Uhrf1^fl/fl^*). (**I**) Total cell numbers of ex-Foxp3 and Treg cells. For mice started on tamoxifen at 4 weeks of age, *n* = 5 (i*Uhrf1^+/+^*) and 7 (i*Uhrf1^fl/fl^*); for mice started on tamoxifen at 8 weeks of age, *n* = 4 (i*Uhrf1^+/+^*) and 7 (i*Uhrf1^fl/fl^*). * *q* < 0.05, ** *p* or *q* < 0.01, † *p* or *q* < 0.001, ‡ *p* < 0.0001, n.s. not significant by two-way ANOVA with Sidak’s *post-hoc* testing for multiple comparisons (B), Mann-Whitney test (C and H), or the two-stage linear step-up procedure of Benjamini, Krieger, and Yekutieli with *Q* = 5% (G and I). For the comparison between genotypes in (B), F(DFn, DFd) = F(1, 42) = 51.4 with *p* < 0.0001.

**Supplemental Figure 5.**
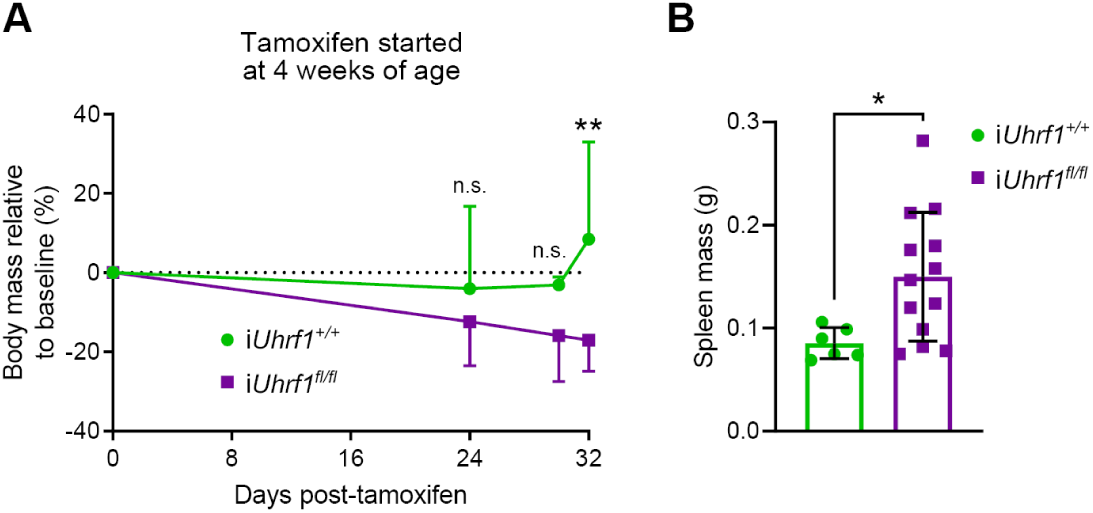
Phenotype of *iUhrf1^fl/fl^* mice after initiation of tamoxifen at 4 weeks of age. (**A**) Body mass curves for *iUhrf1^+/+^* (*n* = 6) and *iUhrf1^fl/fl^* (*n* = 21) mice initiated on tamoxifen at 4 weeks of age. (**B**) Spleen masses for *iUhrf1^+/+^* (*n* = 6) and *iUhrf1^fl/fl^* (*n* = 13) mice after 4 weeks of tamoxifen. Summary plots show all data points with mean and standard deviation. * *p* < 0.05, ** *p* < 0.01, n.s. not significant by two-way ANOVA with Sidak’s *post-hoc* testing for multiple comparisons (A) or Mann-Whitney test (B). For the comparison between genotypes in (A), F(DFn, DFd) = F(1, 59) = 14.6 with *p* = 0.0003.

### Induction of Treg cell-specific Uhrf1 deficiency promotes generation of ex-Foxp3 cells

The i*Uhrf1^fl/fl^* mice permitted tracking the Foxp3^+^ Treg cell lineage to determine whether loss of Uhrf1 promotes generation of ex-Foxp3 cells that would indicate loss of Foxp3^+^ Treg cell identity. Following tamoxifen administration to i*Uhrf1^fl/fl^* mice, cells actively expressing *Foxp3* are labeled with GFP, whereas tdTomato identifies cells that have undergone *Foxp3*-Cre-mediated loss of *Uhrf1* irrespective of active *Foxp3* expression (see Figure 4A). TdTomato also serves as a Foxp3^+^ cell lineage tag in both i*Uhrf1^+/+^* and i*Uhrf1^fl/fl^* mice, marking cells that have undergone *Foxp3*-Cre-mediated recombination of the *ROSA26* locus. Approximately 0.5% of splenic CD3ε^+^CD4^+^ T cells expressed tdTomato in tamoxifen-naïve mice; after 4 weeks of tamoxifen chow, labeling with tdTomato was nearly complete, with only 1% of splenic CD3ε^+^CD4^+^ T cells expressing *Foxp3*-GFP but not the tdTomato label (**Supplemental Figure 6A**). The tdTomato label was highly specific to CD3ε^+^CD4^+^ T cells, with < 0.25% of CD3ε^−^CD4^−^ and CD3ε^+^CD4^−^ cells expressing the tdTomato label in either i*Uhrf1^+/+^* or i*Uhrf1^fl/fl^* mice (**Supplemental Figure 6B**). As reviewed in the Introduction, ex-Foxp3 cells detected in i*Uhrf1^+/+^* mice represent transient Foxp3 induction in a minor population of CD4^+^ T cells that does not represent epigenetic reprogramming of mature Treg cells (26, 27). Thus, in i*Uhrf1^fl/fl^* mice, any ex-Foxp3 cells detected in excess of those observed in control mice represent loss of Foxp3^+^ Treg cell lineage identity.

**Supplemental Figure 6.**
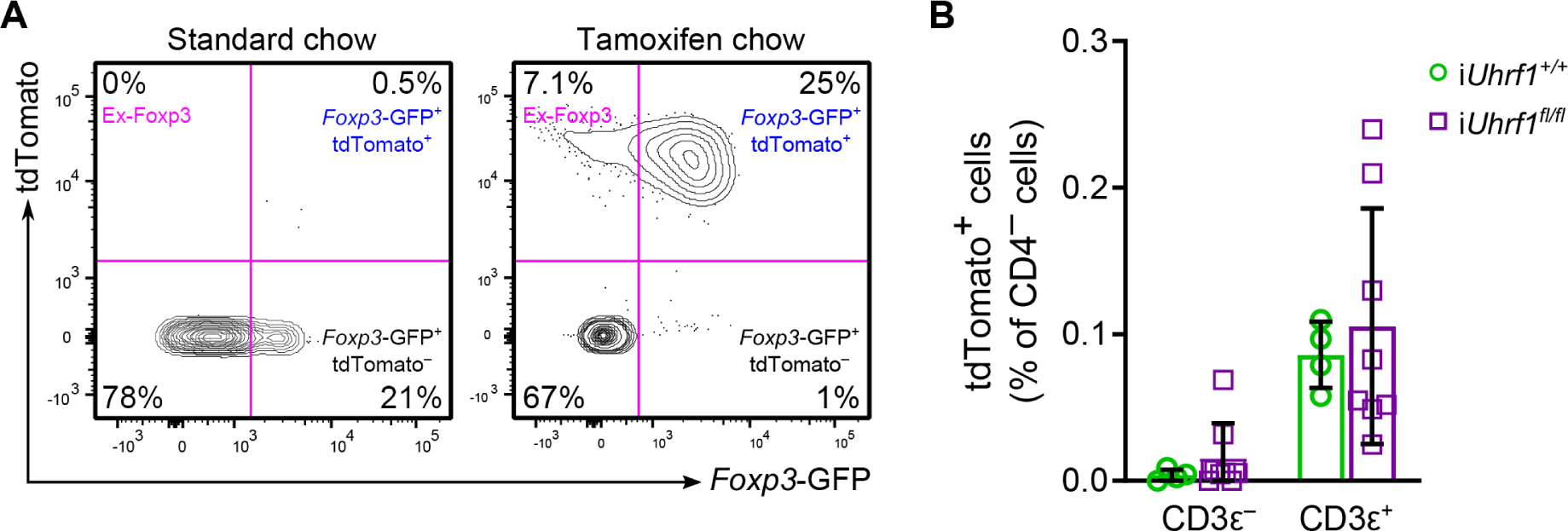
Validation of the Foxp3^+^ lineage-tracing construct. (**A**) 8-week-old i*Uhrf1^+/+^* mice received either standard chow or tamoxifen chow for 4 weeks. Representative contour plots of splenic CD3ε^+^CD4^+^ cells are shown. (**B**) 8-week-old i*Uhrf1^+/+^* and i*Uhrf1^fl/fl^* mice received tamoxifen chow for 4 weeks. Summary plots show all data points with mean and standard deviation as a fraction of splenic CD4^−^ cells. *n* = 4 (*iUhrf1^+/+^*) and 8 (*iUhrf1^fl/fl^*).

Using this system, we found that both 4- and 8-week-old i*Uhrf1^fl/fl^* mice displayed a significant increase in the proportion of ex-Foxp3 (*Foxp3*-GFP^−^tdTomato^+^) cells as a fraction of labeled (tdTomato^+^) cells compared with i*Uhrf1^+/+^* mice (**Figure** 4, F **and** G). The ratio of ex-Foxp3 cells to Treg cells was increased among both age groups, although 4-week-old i*Uhrf1^fl/fl^* mice had a greater increase in the ratio than 8-week-old mice (**Figure 4H**), likely reflecting ongoing thymic Treg cell development at this age. Both 4- and 8-week-old mice exhibited an increase in total ex-Foxp3 cells after 4 weeks of tamoxifen; however, only 4-week-old mice exhibited a concomitant decrease in total Treg cells, again suggesting the deleterious effect of losing Uhrf1 on developing thymic Treg cells prior to stabilization of a robust, self-renewing Treg cell population (**Figure 4I**). Altogether, these data reveal that Uhrf1 is required for stabilization of Treg cell identity and suppressive function, as induced loss of Uhrf1 in Treg cells led to spontaneous inflammation and generation of ex-Foxp3 cells in juvenile and adult mice.

### Ex-Foxp3 cells generated following loss of Uhrf1 exhibit a distinct inflammatory gene expression profile

To define the molecular features of ex-Foxp3 cells resulting from loss of Uhrf1, we used a pulse-chase experimental design in which 8-week-old i*Uhrf1^+/+^* and i*Uhrf1^fl/fl^* mice received tamoxifen for 2 weeks followed by standard chow for 4 weeks (**Figure 5A**). The pulse period allowed tracking of ex-Foxp3 cells; the chase period allowed accumulation of Uhrf1-sufficient (*Foxp3*-GFP^+^tdTomato^−^) cells that were able to suppress the spontaneous systemic inflammation observed in the extended pulse-only experiments (**Supplemental Figure 7, A-E**). Examination of the CD3ε^+^CD4^+^Foxp3^−^ T cell compartment revealed an increased frequency of CD44^hi^CD62L^lo^ cells in the spleens of i*Uhrf1^fl/fl^* mice, consistent with the increased activation of ex-Foxp3 cells observed in i*Uhrf1^fl/fl^* mice as revealed by the transcriptional profiling data below. Principal component analysis of 1,270 differentially expressed genes with an FDR *q*-value < 0.05 revealed distinct clusters based on cell type (*Foxp3*-GFP^+^tdTomato^+^ and ex-Foxp3) and genotype (i*Uhrf1^+/+^* and i*Uhrf1^fl/fl^*) (**Figure 5B**). *K*-means clustering of these differentially expressed genes demonstrated a Uhrf1-dependent signature (top cluster), loss of the core Foxp3-dependent Treg gene signature among ex-Foxp3 cells of both genotypes (middle cluster), and a group of genes significantly upregulated in the ex-Foxp3 cells of i*Uhrf1^fl/fl^* mice (bottom cluster) (**Figure 5C** and **Supplemental Figure 7F**). The bottom cluster contained genes associated with a Th1-skewed effector T cell phenotype, including *Gmza*, *Ifng*, *Il7r*, and the master regulator of the Th1 lineage, *Tbx21*. These transcriptional profiles illuminate a gene expression signature characterized by increased activation and Th1 skewing of ex-Foxp3 cells generated following loss of Uhrf1.

**Figure 5.**
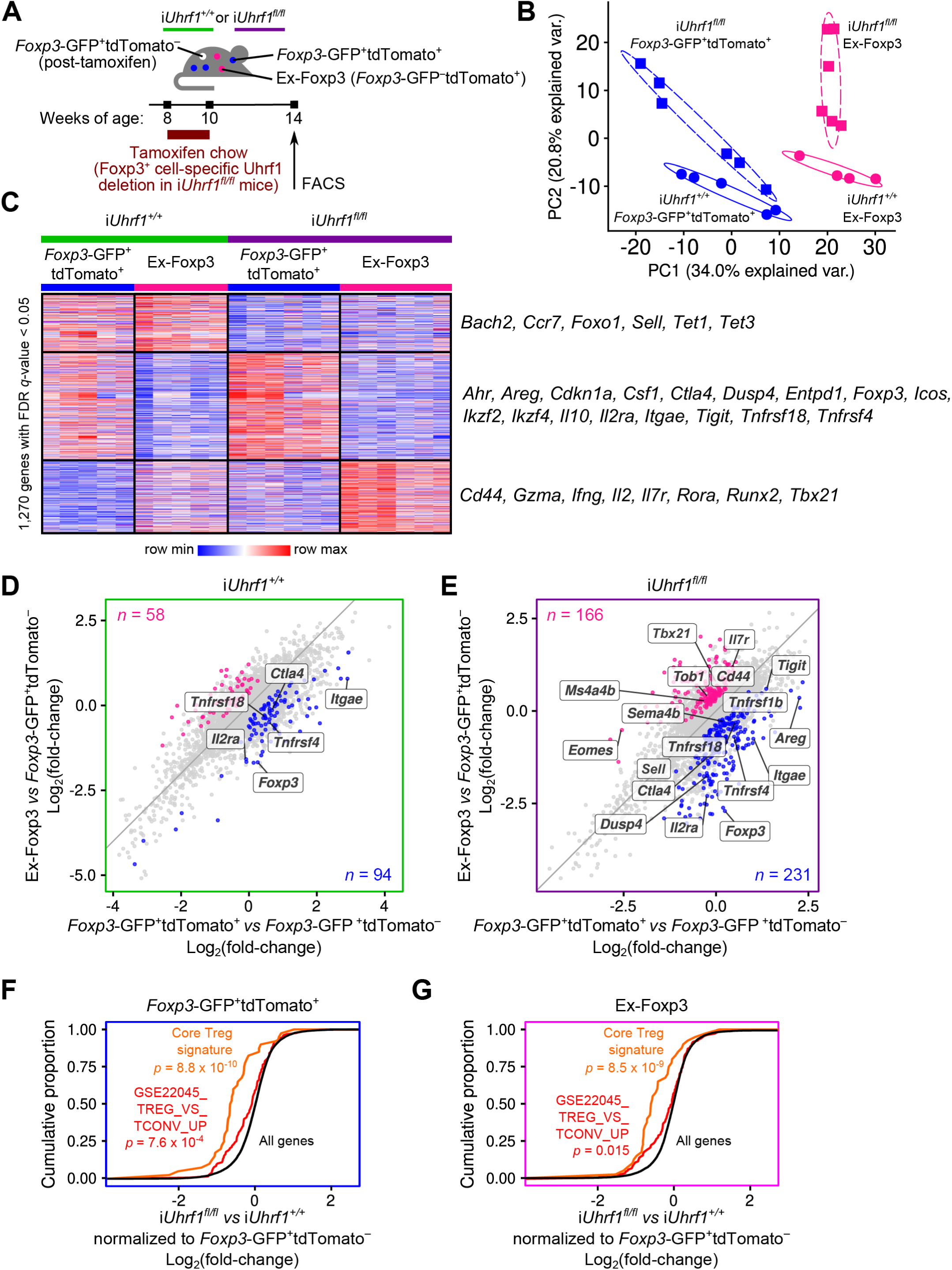
Induction of Uhrf1 deficiency generates ex-Foxp3 cells with distinct inflammatory transcriptional programs. (**A**) Schematic of the pulse-chase experimental design. (**B**) Principal component analysis of 1,270 differentially expressed genes identified from a generalized linear model and ANOVA-like testing with FDR q-value < 0.05. Ellipses represent normal contour lines with one standard deviation probability. (**C**) *K*-means clustering of 1,270 genes with an FDR *q*-value < 0.05 comparing the cell populations from (B) with *k* = 3 and scaled as *z*-score across rows. Genes of interest are annotated. (**D and E**) Fold-change-fold-change plots for i*Uhrf1^+/+^* (D) and i*Uhrf1^fl/fl^* (E) mice of *Foxp3*-GFP^+^tdTomato^+^ versus *Foxp3*-GFP^+^tdTomato^−^ and ex-Foxp3 versus *Foxp3*-GFP^+^tdTomato^−^ highlighting genes exhibiting an increase (*q* < 0.05) in *Foxp3*-GFP^+^tdTomato^+^ versus ex-Foxp3 (blue dots) and ex-Foxp3 versus *Foxp3*-GFP^+^tdTomato^+^ (pink dots). Numbers of differentially expressed genes are indicated. Genes of interest are annotated. (**F**) Cumulative distribution function plots comparing the *Foxp3*-GFP^+^tdTomato^+^ cells from i*Uhrf1^+/+^* versus i*Uhrf1^fl/fl^* mice with each population normalized to the *Foxp3*-GFP^+^tdTomato^−^ population sorted from their respective genotype. The cumulative proportion of all genes (black), a Treg cell-defining gene set (40) (orange), and the GSE22045_TREG_VS_TCONV_UP gene set (43) (red) are shown. (G) Cumulative distribution function plots as in (F) comparing the ex-Foxp3 cells from i*Uhrf1^+/+^* versus i*Uhrf1^fl/fl^* mice. *n* = 5 (i*Uhrf1^+/+^*) and 6 (i*Uhrf1^fl/fl^*). The *p*-values resulting from Kolmogorov-Smirnov tests for cumulative distributions comparing all genes against either gene set are shown in (F and G). FACS, fluorescence-activated cell sorting.

**Supplemental Figure 7.**
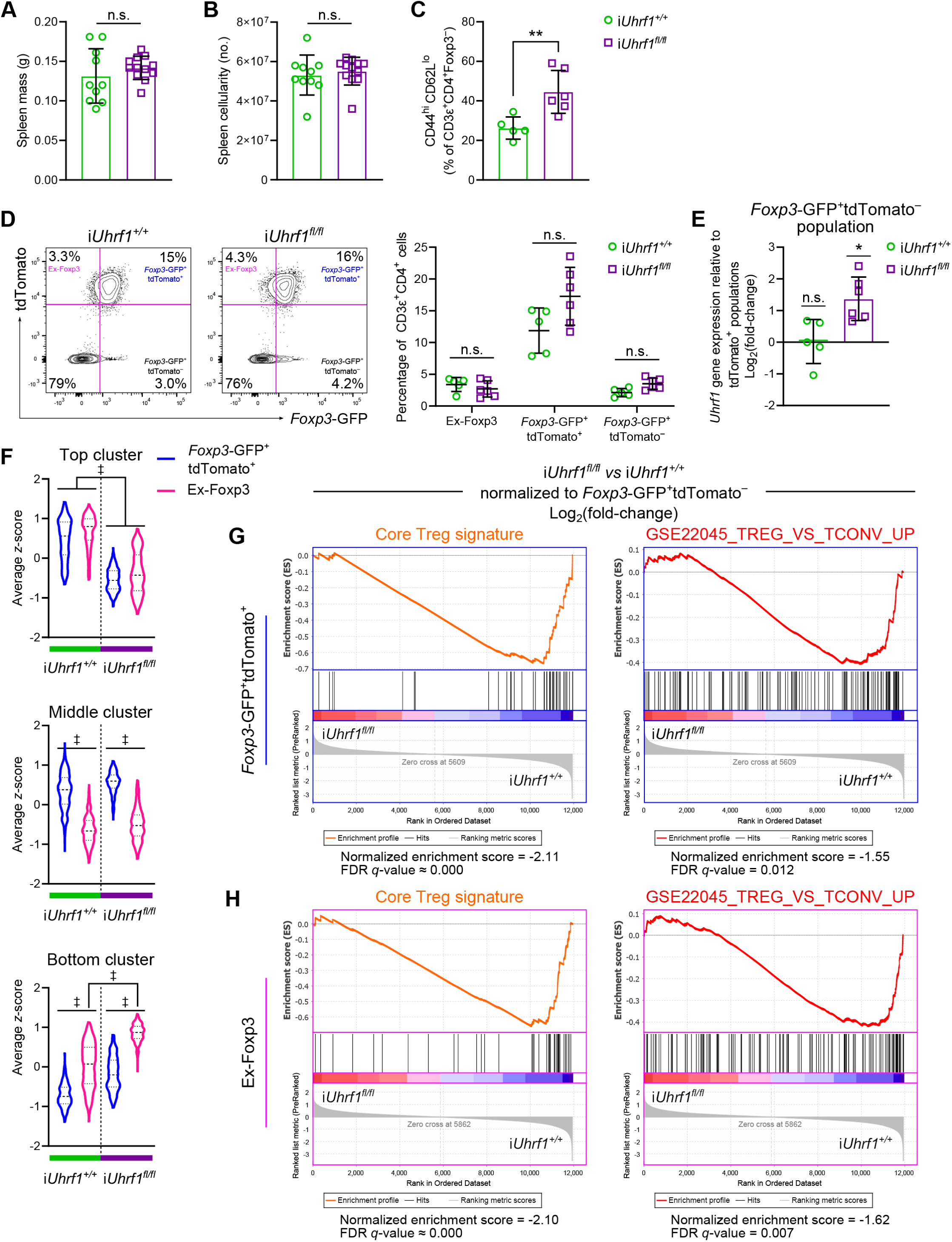
Extended phenotypic and transcriptional analysis of the pulse-chase model. (**A**) Splenic mass of *iUhrf1^+/+^* (*n* = 10) and *iUhrf1^fl/fl^* (*n* = 12) mice after a 2-week pulse of tamoxifen chow followed by 4 weeks of standard chow. (**B**) Spleen cellularity of *iUhrf1^+/+^* (*n* = 10) and *iUhrf1^fl/fl^* (*n* = 12) mice. (**C**) Activation state of splenic CD3ε^+^CD4^+^Foxp3^−^ cells. *n* = 5 (*iUhrf1^+/+^*) and 6 (*iUhrf1^fl/fl^*). (**D**) Representative contour plots of splenic live CD3ε^+^CD4^+^ cells are shown with accompanying summary data. *n* = 5 (*iUhrf1^+/+^*) and 6 (*iUhrf1^fl/fl^*). (**E**) *Uhrf1* gene expression in *Foxp3*-GFP^+^tdTomato^−^ cells relative to the tdTomato^+^ populations. *n* = 5 (*iUhrf1^+/+^*) and 6 (*iUhrf1^fl/fl^*). (**F**) Average *z*-scores for the top, middle, and bottom *k*-means clusters shown in Figure 5C. (**G** and **H**) Gene set enrichment plots testing for enrichment of a Treg cell-defining gene set (40) or the GSE22045_TREG_VS_TCONV_UP gene set (43) with genes ordered by log2(fold-change) in average expression comparing *Foxp3*-GFP^+^tdTomato^+^ (G) or ex-Foxp3 (H) cells from *iUhrf1^fl/fl^* versus *iUhrf1^+/+^* mice. Expression values are normalized to the *Foxp3*-GFP^+^tdTomato^−^ population sorted from the respective genotype. The normalized enrichment scores (NES) and FDR *q*-values associated with these tests are shown. Summary plots show all data points with mean and standard deviation; violin plots show median and quartiles. *n* = 5 (i*Uhrf1^+/+^*) and 6 (i*Uhrf1^fl/fl^*). * *p* < 0.05, ** *p* < 0.01, ‡ *q* < 0.0001, n.s. not significant by Mann-Whitney test (A-C), the two-stage linear step-up procedure of Benjamini, Krieger, and Yekutieli with *Q* = 5% (D), one-way ANOVA (E), or a mixed-effects analysis with the two-stage linear step-up procedure of Benjamini, Krieger, and Yekutieli with *Q* = 5% (F). For the comparisons in (E) between i*Uhrf1^+/+^* cell populations, F(DFn, DFd) = F(2, 12) = 1.4 with *p* = 0.28; for i*Uhrf1^fl/fl^* cell populations, F(DFn, DFd) = F(2, 15) = 4.3 with *p* = 0.03.

In contrast with the extended pulse experiment shown in Figure 4, the frequency of each labeled population was similar between i*Uhrf1^+/+^* and i*Uhrf1^fl/fl^* mice following the pulse-chase period. Thus, we used the Uhrf1-sufficient *Foxp3*-GFP^+^tdTomato^−^ cell population as an internal control for the transcriptional analysis of both i*Uhrf1^+/+^* and i*Uhrf1^fl/fl^* mice (see Supplemental Figure 7D). After normalizing to this Uhrf1-sufficient cell population within each genotype, the ex-Foxp3 cells of i*Uhrf1^+/+^* mice exhibited deficiency of the core Foxp3-dependent Treg cell signature (**Figure 5D**). The ex-Foxp3 cells of i*Uhrf1^fl/fl^* mice also displayed loss of this core signature but simultaneously upregulated the expression of genes classically associated with impaired Treg cell function and gain of conventional effector T cell function, including *Tbx21*, *Eomes, Ms4a4b, Tob1*, and *Il7r* in addition to evidence of activation (upregulation of *Cd44* and downregulation of *Sell*) (**Figure 5E**). Consistent with the results of the Uhrf1 chimeric knockout experiments shown in Figure 3, examination of *Foxp3*-GFP^+^tdTomato^+^ cells from i*Uhrf1^fl/fl^* versus i*Uhrf1^+/+^* mice with respect to previously annotated gene sets of core Treg cell identity (40) and Treg versus conventional T cell profiles (43) revealed loss of the core Treg cell signature and a shift toward a conventional effector T cell state (**Figure 5F** and **Supplemental Figure 7G**). Even when comparing ex-Foxp3 cells between genotypes and normalizing to the internal Uhrf1-sufficient *Foxp3*-GFP^+^tdTomato^−^ cell population, i*Uhrf1^fl/fl^* mice still exhibited a greater loss of Treg cell identity and skewing toward a conventional effector T cell state compared with i*Uhrf1^+/+^* mice (**Figure 5G** and **Supplemental Figure 7H**). Collectively, these unsupervised analyses demonstrate that loss of Uhrf1 results in generation of ex-Foxp3 cells displaying an excessively activated and inflammatory profile.

### Altered DNA methylation patterns underlie the gain of inflammatory signature and loss of Treg cell signature observed in Uhrf1-deficient ex-Foxp3 cells

We performed genome-wide CpG methylation profiling on the sorted *Foxp3*-GFP^+^tdTomato^+^ and ex-Foxp3 cells from i*Uhrf1^+/+^* and i*Uhrf1^fl/fl^* mice obtained following the pulse-chase experiment illustrated in Figure 5A. Principal component analysis of differentially methylated cytosines revealed nominal differences between the *Foxp3*-GFP^+^tdTomato^+^ and ex-Foxp3 cells from i*Uhrf1^+/+^* mice but substantial spread between the same cell types obtained from i*Uhrf1^fl/fl^* mice (**Figure 6A**). Similar to the findings from Uhrf1 chimeric knockout animals, *Foxp3*-GFP^+^tdTomato^+^ cells from i*Uhrf1^fl/fl^* mice contained a global hypomethylation pattern that was further hypomethylated in ex-Foxp3 cells (**Figure 6B**). In contrast, Uhrf1-sufficient ex-Foxp3 cells exhibited only a minor degree of hypomethylation relative to *Foxp3*-GFP^+^tdTomato^+^ cells. To explore these differential methylation patterns in more detail, we performed an unsupervised procedure that revealed 17,249 DMRs that were within 5 kb of 8,101 gene bodies (inclusive). These putative regulatory elements did not exhibit differential methylation between *Foxp3*-GFP^+^tdTomato^+^ and ex-Foxp3 cells from i*Uhrf1^+/+^* mice; however, both cell types were hypomethylated in i*Uhrf1^fl/fl^* mice, which displayed further reductions in methylation within ex-Foxp3 compared with *Foxp3*-GFP^+^tdTomato^+^ cells (**Figure 6C**). These DMRs were located near 642 differentially expressed genes (a majority of 1,270), a finding that was unlikely due to chance alone (**Figure 6D**).

**Figure 6.**
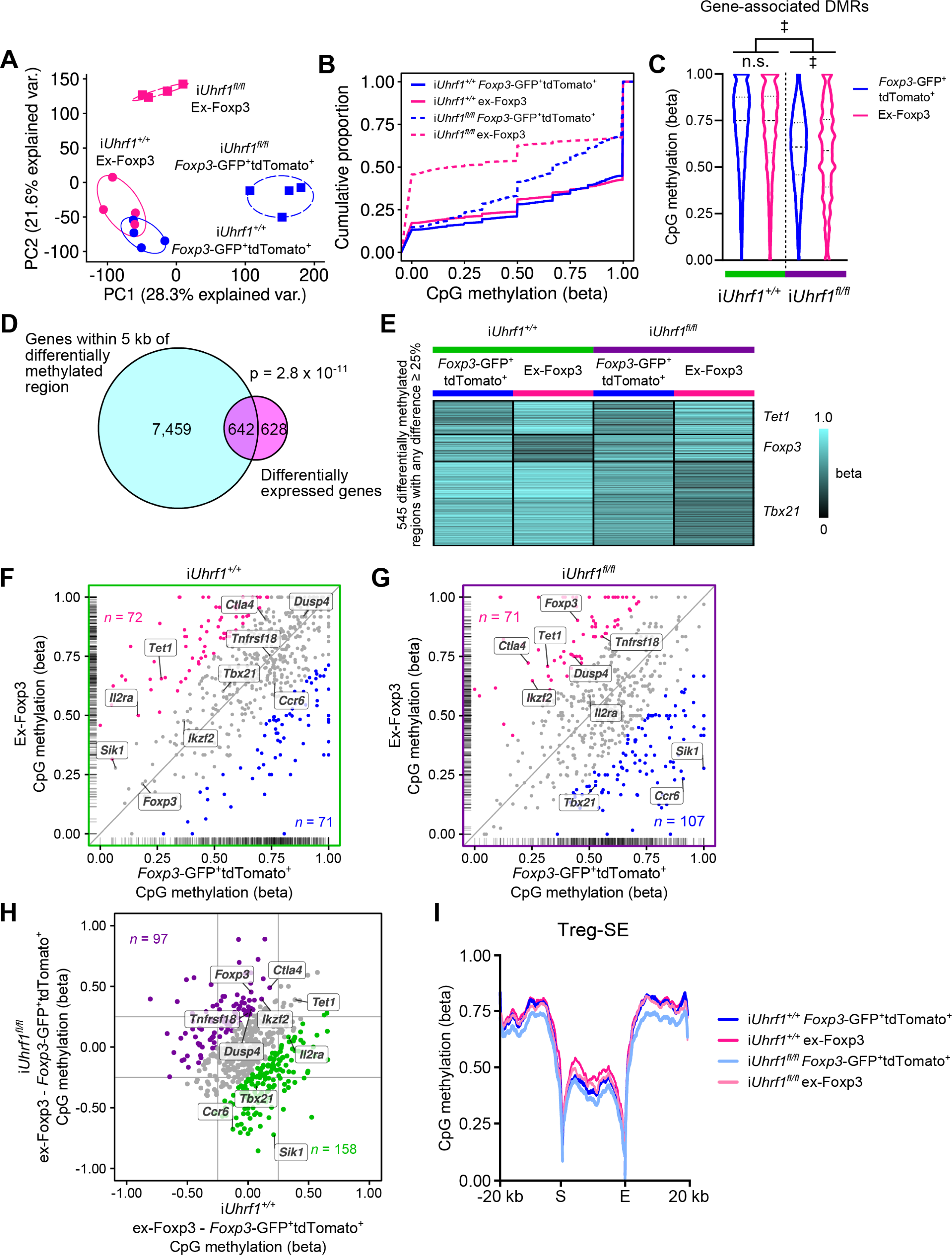
Altered DNA methylation patterns explain the transcriptional reprogramming of Uhrf1-deficient Treg and ex-Foxp3 cells. (**A**) Principal component analysis of 202,525 differentially methylated CpGs identified from a beta-binomial regression model with an arcsine link function fitted using the generalized least square method and Wald-test FDR *q*-value < 0.05. Ellipses represent normal contour lines with one standard deviation probability. (**B**) Cumulative distribution function plot of differentially methylated CpGs. (**C**) CpG methylation at 17,249 differentially methylated regions (DMRs) within 5 kb of gene bodies (inclusive). (**D**) Venn diagram of genes associated with a DMR and differentially expressed genes. (**E**) *K*-means clustering of 545 DMRs with a difference > 25% between any groups and an FDR q-value < 0.05 that were grouped and then averaged by unique gene association. *K* = 3 with scaling as beta scores across rows. (**F and G**) Scatter plots comparing DMRs within *Foxp3*-GFP^+^tdTomato^+^ cells versus ex-Foxp3 cells from i*Uhrf1^+/+^* mice (F) and i*Uhrf1^fl/fl^* mice (G). Interior axis ticks (rug plots) represent positions of DMRs on each axis. Colored points represent > 25% difference between ex-Foxp3 cells (pink) and *Foxp3*-GFP^+^tdTomato^+^ cells (blue) with the number of DMRs in each such subset shown. (**H**) Difference-difference plot comparing the difference in DMR methylation status between ex-Foxp3 and *Foxp3*-GFP^+^tdTomato^+^ cells in i*Uhrf1^+/+^* mice and i*Uhrf1^fl/fl^* mice. Colored points represent > 25% difference between i*Uhrf1^+/+^* mice (green) and i*Uhrf1^fl/fl^* mice (purple) with the number of DMRs in each such subset shown. (**I**) Metagene analysis of CpG methylation across the start (S) through the end (E) of Treg cell-specific super-enhancer elements (Treg-SE) as defined in reference (19). Violin plots show median and quartiles. *n* = 4 mice per cell type for both genotypes. A hypergeometric *p*-value is shown in (D). ‡ *q* < 0.0001, n.s. not significant by a mixed-effects analysis with the two-stage linear step-up procedure of Benjamini, Krieger, and Yekutieli with *Q* = 5% (C).

We next examined DMRs found near differentially expressed genes and selected those regions with greater than 25% difference between any groups. Functional enrichment analysis of these regions using the Genomic Regions Enrichment of Annotations Tool (GREAT) (44) and the Mouse Genome Informatics Phenotype ontology (45) revealed significant enrichment in phenotypes that are characterized by abnormal adaptive immunity and dysregulated T cell development and physiology (**Supplemental Figure 8, A and B**). We then averaged the methylation level of DMRs associated with unique genes and defined three *k*-means clusters (**Figure 6E**); the cluster structure mirrored the *k-*means transcriptional analysis in Figure 5C. Inspection of the three clusters revealed a large hypomethylated cluster peculiar to Uhrf1-deficient ex-Foxp3 cells that contained the master regulator of the Th1 lineage, *Tbx21*. Indeed, the DMR(s) associated with *Tbx21* and other inflammatory genes were hypomethylated in Uhrf1-deficient but not Uhrf1-sufficient ex-Foxp3 cells (**Figure 6, F-H and Supplemental Figure 8C**). Interestingly, DMRs near core Treg cell signature genes, including *Foxp3*, became methylated in Uhrf1-deficient but not Uhrf1-sufficient ex-Foxp3 cells (**Supplemental Figure 8D**). Because the primary mechanistic effect of Uhrf1 loss is *hypo*methylation, the observed *hyper*methylation of Treg cell signature genes appears to represent a secondary effect of derepressing the inflammatory program. Compared with the other populations, methylation across Treg-SE was greatest within the ex-Foxp3 cells of i*Uhrf1^+/+^* mice, likely reflecting their developmental origin as unstable Foxp3^+^ or potential Treg cells that transiently expressed Foxp3 after thymic emigration (28) (**Figure 6I**). Altogether, genome-wide DNA methylation profiling and an unsupervised analytic approach revealed alterations in DNA methylation patterning that explained the transcriptional and phenotypic reprogramming that results from loss of Uhrf1.

**Supplemental Figure 8.**
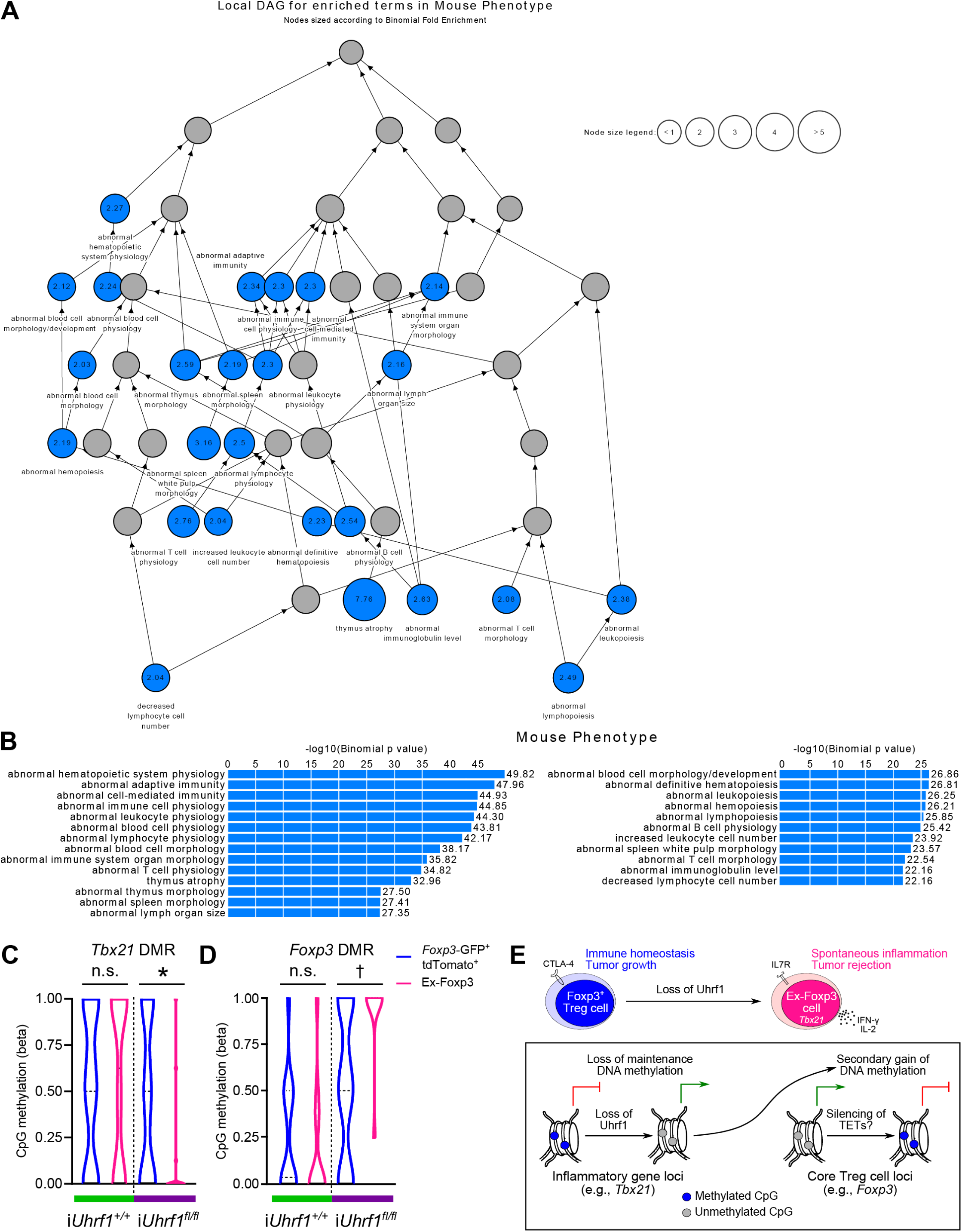
Extended DNA methylation analysis of the pulse-chase model. (**A**) Directed acyclic graph (DAG) based on the enriched terms (shown in blue) from the Mouse Genome Informatics Phenotype ontology (45). Nodes are sized according to binomial fold enrichment. (**B**) Bar chart of binomial *p*-values based on the analysis shown in (A). (**C and D**) CpG methylation at the differentially methylated region (DMR) located at the *Tbx21* locus (C, chr11:97,112,193-97,113,285) and the *Foxp3* locus (D, chrX:7,580,236-7,581,944). (**E**) Graphical abstract of the proposed model. Violin plots show median and quartiles. *n* = 4 mice per cell type for both genotypes. * *q* < 0.05, † *q* < 0.001, n.s. not significant by a mixed-effects analysis with the two-stage linear step-up procedure of Benjamini, Krieger, and Yekutieli with *Q* = 5% (C and D).

### Loss of Uhrf1 in mature Treg cells results in enhanced tumor immunity and accumulation of intra-tumoral ex-Foxp3 cells

To further test the effect of induced Uhrf1 loss on Treg cell suppressive function and ex-Foxp3 cell effector function, we evaluated the ability of i*Uhrf1^fl/fl^* mice to grow B16 melanoma tumors (46). Compared with i*Uhrf1^+/+^* controls, i*Uhrf1^fl/fl^* mice exhibited slowed tumor growth and the beginning of tumor regression at 3 weeks post-injection—the time when control mice required euthanasia due to tumor ulceration (**Figure 7A**). When assessed *ex vivo* at 21 days post-injection, tumor volumes and masses were significantly lower among i*Uhrf1^fl/fl^* mice compared with control mice (**Figure 7**, B **and** C). Flow cytometry analysis revealed a substantial increase in the frequency of ex-Foxp3 cells within the tumors of i*Uhrf1^fl/fl^* mice, which also exhibited a paucity of Treg cells (**Figure 7D**). These experiments illustrate that mature Treg cells require Uhrf1 to suppress host-versus-tumor immunity and that induction of Treg cell-specific Uhrf1 deficiency promotes generation of ex-Foxp3 cells during the immune response to malignancy.

**Figure 7.**
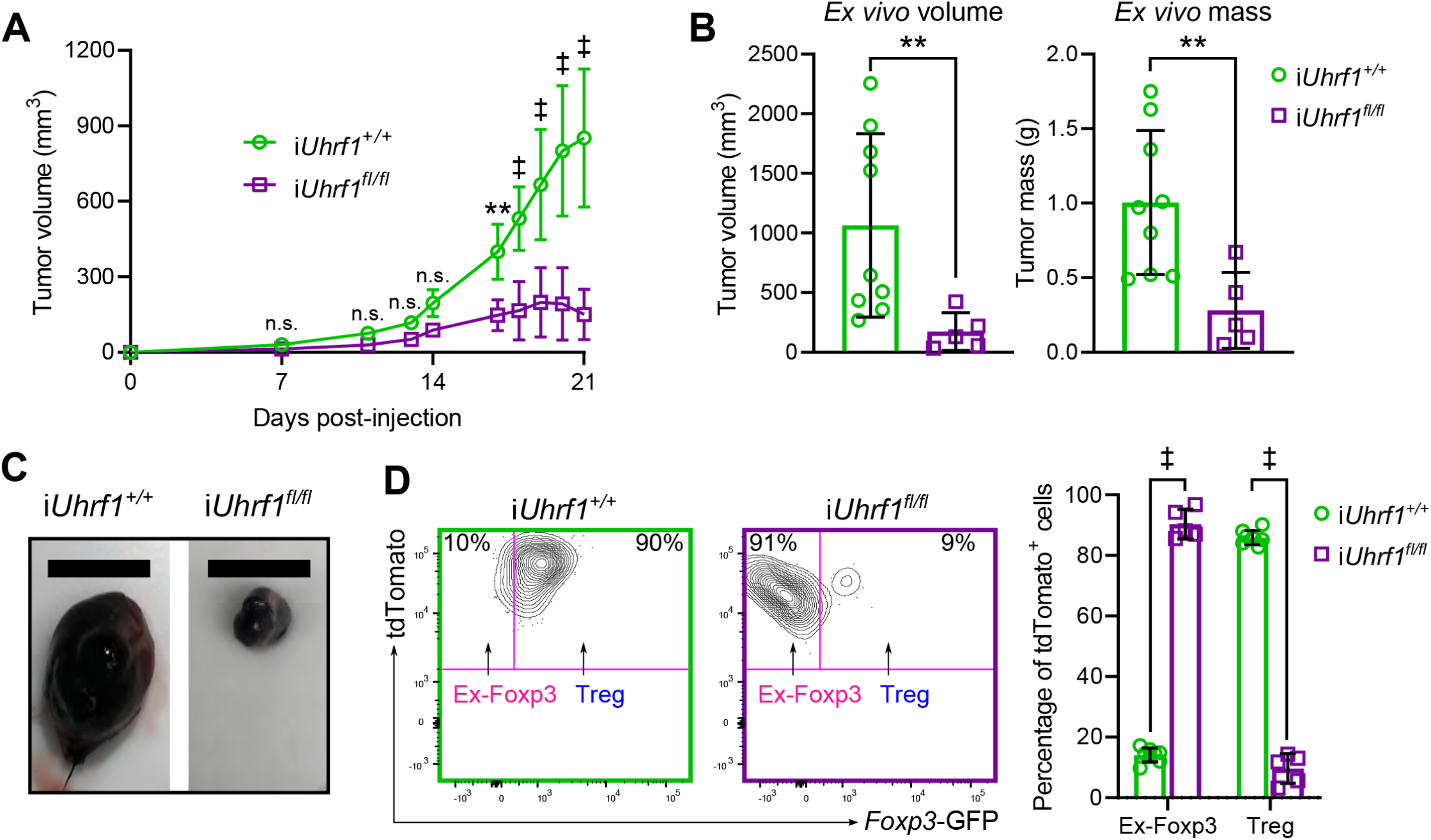
Induced loss of Uhrf1 in Treg cells inhibits tumor growth and promotes infiltration of intra-tumoral ex-Foxp3 cells. (**A**) Growth of B16 melanoma cells in i*Uhrf1^+/+^* and i*Uhrf1^fl/fl^* mice. Tamoxifen chow was administered to 8-week-old mice beginning 3 weeks before tumor cell injection. (**B**) *Ex vivo* tumor volumes and masses measured after post-mortem tumor resection on day 21 post-injection. (**C**) Photomicrographs of resected tumors. Scale bar represents 1 cm. Tumor experiments were performed over two separate experiments with total *n* = 9 (i*Uhrf1^+/+^*) and 5 (i*Uhrf1^fl/fl^*). (**D**) Representative flow cytometry contour plots gated on splenic CD3ε^+^CD4^+^tdTomato^+^ cells showing the percentage of intra-tumoral ex-Foxp3 (*Foxp3*-GFP^−^) and Treg (*Foxp3*-GFP^+^) cells, summarized in the accompanying graph. *n* = 8 (i*Uhrf1^+/+^*) and 5 (i*Uhrf1^fl/fl^*). Summary plots show all data points with mean and standard deviation. ** *p* < 0.01, ‡ *p* or *q* < 0.0001, n.s. not significant by two-way ANOVA with Sidak’s *post-hoc* testing for multiple comparisons (A), Mann-Whitney test (B), or the two-stage linear step-up procedure of Benjamini, Krieger, and Yekutieli with *Q* = 5% (D). For the comparison between genotypes in (A), F(DFn, DFd) = F(1, 120) = 130.0 with *p* < 0.0001.

Collectively, our findings demonstrate that Foxp3^+^ Treg cells require Uhrf1-mediated maintenance of DNA methylation at inflammatory gene loci for their development and functional stability (**Supplemental Figure 8E**).

## Discussion

Regulatory T cells develop from the complex interplay of *trans*- and *cis*-regulatory mechanisms and are maintained as a stable, self-renewing population (17, 19, 26, 29, 47). Here, we determined that Treg cells require maintenance DNA methylation mediated by the epigenetic regulator Uhrf1 for establishment of the Foxp3^+^ Treg cell lineage as well as stability of Treg cell identity and suppressive function. Loss of Uhrf1 at the Foxp3^+^ stage of thymic Treg cell development resulted in Treg cell deficiency, leading to a scurfy-like lethal inflammatory disorder. Studies using non-inflamed Uhrf1 chimeric knockout animals revealed the cell-autonomous and Foxp3-independent requirement for Uhrf1-mediated DNA methylation in stabilizing the lineage by repressing effector T cell transcriptional programs. Induced deficiency of Uhrf1 in mature Treg cells was sufficient to derepress loci encoding inflammatory genes, resulting in loss of Treg cell identity and suppressive function, generation of inflammatory ex-Foxp3 cells, spontaneous inflammation, and augmented tumor immunity. Taken together, these results demonstrate the necessity of maintenance DNA methylation in stabilizing Treg cell identity and suppressive function during both development and self-renewal of the lineage. Our findings change the existing model of DNA hypomethylation as the dominant feature of the Treg cell epigenome by demonstrating an essential role for methylation-mediated silencing of effector programs in the development and stability of the Treg cell lineage.

In our studies, we found that Uhrf1 serves as an essential regulator that is required to maintain DNA methylation at defined genomic loci during both Treg cell development and self-renewal. Unlike *Uhrf1^fl/fl^Cd4^Cre^* mice with pan-T cell Uhrf1 deficiency that develop inflammation localized to the colon (37), *Uhrf1^fl/fl^Foxp3^Cre^* mice in our study spontaneously developed widespread inflammation (the scurfy phenotype), indicating a fundamental role for Uhrf1-mediated maintenance of DNA methylation in stabilizing Treg cell identity after Foxp3 induction in the thymus. These findings also suggest that Uhrf1 is required to maintain a specific DNA methylation landscape in pre-Treg (pre-Foxp3) thymocytes that differs from the landscape required after induction of Foxp3 expression. Following thymic emigration, the Treg cell lineage is remarkably stable with self-renewal across the lifespan (29). Indeed, the ex-Foxp3 cells that we observed in adult control animals did not display an inflammatory gene expression profile but did exhibit increased super-enhancer methylation compared with Treg cells, reflecting their developmental origin as either unstable Foxp3^+^ cells or potential Treg cells as defined by Ohkura et al (28). In contrast, the increased number of ex-Foxp3 cells generated following loss of Uhrf1 expressed a Th1-like profile, indicating destabilization of the Treg cell lineage upon loss of maintenance DNA methylation.

We propose a model in which Treg cells require maintenance of DNA methylation at inflammatory gene loci for their development and stability (see Supplemental Figure 8E). In our model, loss of these marks results in derepression of inflammatory genes, including *Tbx21*. A secondary wave of methylation then represses core Treg cell loci, including *Foxp3*, to generate inflammatory ex-Foxp3 cells that contribute to spontaneous inflammation and tumor rejection. The genome-wide transcriptional and DNA methylation profiles of Uhrf1-deficient ex-Foxp3 cells support this mechanism, with hypomethylation of the *Tbx21* locus and secondary methylation of the *Foxp3* locus. How gain of the effector program results in silencing of *Foxp3* and other Treg cell-defining loci remains an open question. Our data support the speculation that methylation-mediated silencing of TET enzyme expression in Uhrf1-deficient ex-Foxp3 cells contributes to gain of methylation at core Treg cell loci, ultimately downregulating their expression (48).

Our results inform the rational design of DNA methylating or demethylating agents for the purpose of immunomodulatory therapy. Numerous lines of evidence demonstrate that pharmacologic inhibition of DNA methyltransferase activity induces Foxp3 expression in CD4^+^ T cells and promotes Treg cell suppressive function in adult mice (20-22, 49-51). Based on these studies, we initially hypothesized that loss of Uhrf1 in adult mice would promote Treg cell generation and augment their suppressive function. Nevertheless, our results show that adult mice with induced loss of Uhrf1 spontaneously develop widespread inflammation reminiscent of the developmental phenotype observed in the constitutive conditional knockout animals, scurfy mice, and humans with the autoimmune IPEX syndrome. Indeed, missense variants of *UHRF1BP* (Uhrf1 binding protein) are associated with the development of systemic lupus erythematosus (52). Thus, our results suggest caution when applying pharmacologic approaches to epigenetic modification in immune-dysregulated conditions such as autoimmune and malignant disorders.

Collectively, our findings reveal that Foxp3^+^ Treg cells require maintenance DNA methylation mediated by the epigenetic regulator Uhrf1 for both development and stability of the lineage. Loss of Uhrf1-mediated DNA methylation during both development and self-renewal is sufficient to disrupt the lineage, generate inflammatory ex-Foxp3 cells, and cause inflammatory pathology and tumor rejection. Because of the powerful role that DNA methylation programming plays in determining the stability of the Treg cell lineage, pharmacologic stabilization or disruption of the DNA methylation maintenance function of Uhrf1 could be explored to address inflammatory or malignant disorders, respectively.

## Methods

### Mice

C57BL/6 *Uhrf1^fl/fl^* mice, which harbor *loxP* sequences flanking exon 4 (ENSMUSE00000139530) of the *Uhrf1* gene (ENSMUSG00000001228), were a kind gift from the laboratory of Srinivasan Yegnasubramanian, MD, PhD (Johns Hopkins University). *Foxp3^YFP-Cre^* (stock no. 016959), *Foxp3^eGFP-CreERT2^* (stock no. 016961), and *ROSA26Sor^CAG-tdTomato^* (Ai14, stock no. 007908) mice were obtained from The Jackson Laboratory. All animals were genotyped using services provided by Transnetyx, Inc. with primers provided by The Jackson Laboratory or reported below. Experimental randomization and blinding were not performed due to obvious external phenotype; however, littermate controls were used whenever possible. Animals received water *ad libitum*, were housed at a temperature range of 20-23 °C under 14/10-hour light/dark cycles, and received standard rodent chow except for animals that received chow containing tamoxifen (Envigo, 500 mg/kg of chow ≈ daily dose of 40-80 mg/kg) as noted. All animal experiments and procedures were conducted in accordance with the standards established by the United States Animal Welfare Act set forth in National Institutes of Health guidelines and were approved by the Institutional Animal Care and Use Committee (IACUC) at Northwestern University.

### Tissue and cell preparation

Organs were prepared for histological assessment with sectioning and hematoxylin and eosin staining as previously described (20, 46). For preparation of colonic tissue for flow cytometry, colons were collected from mice, flushed with 10 mL ice-cold PBS, and washed twice with RT HBSS + 2% fetal bovine serum (FBS). Colons were then cut into 1-cm pieces, placed in a 50-mL conical tube containing 10 mL RT HBSS + 2% FBS + 2 mM EDTA, and incubated for 15 minutes at 37 °C with rotation. Supernatant was discarded and the colon was washed twice with HBSS, cut into 2-mm pieces, placed in 10 mL of pre-warmed RPMI + 10% FBS + 1.5 mg/mL Collagenase D (Roche 11088866001) + 0.05 mg/mL Collagenase V (Sigma C9263) and incubated for 40 minutes with rotation. Digested colons were filtered through a 40-µm strainer into 10 mL of MACS buffer (Miltenyi 130-091-221), centrifuged 10 minutes at 1,500 RPM, and resuspended in MACS buffer. Histological evaluations were performed in consultation with a veterinary pathologist through core services provided by the Northwestern University Mouse Histology and Phenotyping Laboratory.

### Flow cytometry analysis and fluorescence-activated cell sorting

Single-cell suspensions of visceral organs and blood were prepared and stained for flow cytometry analysis and fluorescence-activated cell sorting as previously described (20, 46, 53–55) using the reagents and cytometer setup shown in **Supplemental Table 1**. Cell counts of single-cell suspensions were obtained using a hemocytometer with trypan blue exclusion or a Cellometer with AO/PI staining (Nexcelom Bioscience) before preparation for flow cytometry. Data acquisition for analysis was performed using a BD LSRFortessa or Symphony A5 instrument with FACSDiva software (BD). Cell sorting was performed sing the 4-way purity setting on BD FACSAria SORP instruments with FACSDiva software. Fluorescence-minus-one controls were used. Analysis was performed with FlowJo v10.6.1 software. Dead cells were excluded using a live-dead marker for analysis and sorting.

### Cytokine measurements

Following lysis of erythrocytes, single-cell suspensions of spleen and lung tissue and blood at a concentration of 5 x 10^6^ cells/mL were treated with RPMI plus 5 µL/mL Leukocyte Activation Cocktail with GolgiPlug (BD) or RMPI only and incubated for 4 hours at 37 °C. After incubation, cells were spun down, washed, and resuspended in a viability dye (Fixable Viability Dye eFluor 506, eBioscience) for 30 minutes at 4 °C and fixed and permeabilized with the Foxp3/Transcription Factor Staining Buffer Set (eBioscience). Cells were then stained with an antibody cocktail (Fc Block, Foxp3-PE-Cy7, CD4-PerCP-Cy5.5, IL-17A-PE, IFN-γ-PE-CF594, IL-4-APC; see Supplemental Table 1 for details) for 30 minutes at room temperature, washed, and resuspended in PBS with 0.5% bovine serum albumin (BSA) before flow cytometry analysis as above.

### Annexin staining

Single-cell suspensions were incubated with a UV-excitable viability dye (Molecular Probes #L23105) followed by surface staining as above and previously described (20, 46, 53, 54). Cells were then washed and resuspended in annexin-binding buffer (10 mM HEPES, 140 mM NaCl, and 2.4 mM CaCl2; pH 7.4) at a concentration of 1 x 10^6^ cells/mL. 5 µL Annexin V Pacific Blue conjugate (Invitrogen #A35122) per 100 µL of cell suspension was then added and incubated for 15 minutes at room temperature and then washed with annexin-binding buffer before resuspension in PBS with 0.5% BSA for flow cytometry analysis as above.

Uhrf1 *PCR*. DNA was extracted from sorted cells using the Qiagen AllPrep DNA/RNA Micro Kit and quantified using a Qubit 3.0 fluorometer. PCR was performed using the following primers (IDT, standard desalting): Forward (F) CTTGATCTGTGCCCTGCAT, Reverse 1 (R1) ACCTCTGCTCTGATGGCTGT, and Reverse 2 (R2) CCGAGGACACTCAAGAGAGC. As shown in Supplemental Figure 1A, the F and R1 primers flank the 5’ *loxP* site and produce a 198-bp band in WT mice and a 398-bp band in conditional (floxed) mice. The R2 primer sits downstream of the 3’ *loxP* site, resulting in a 247-bp band in cells that have undergone Cre-mediated excision of the floxed sequence. 0.1 ng of template DNA was combined with 10 µL iQ supermix (Bio-Rad) and 2 µL each of 20 µM F primer, 10 µM R1 primer, and 10 µM R2 primer in a total reaction volume of 20 µL. A C1000 Touch Thermal Cycler (Bio-Rad) was programmed for 1 minute at 95 °C; then 15 seconds at 95 °C, 15 seconds at 65 °C, and 30 seconds at 72 °C for 35 cycles; and ending with 7 minutes at 72 °C. 20 µL of PCR product was combined with 32 µL of Magbio High-Prep PCR beads (1.6X clean-up). After a 5 minute incubation, the tubes were placed on a magnet until the solution cleared. The supernatant was removed, the beads were washed twice with 80% ethanol, and the product was eluted in 20 µL water. The cleaned PCR product was run on an Agilent TapeStation 4200 using High Sensitivity D1000 ScreenTape and visualized using TapeStation Analysis Software A.02.01 SR1.

### RNA-sequencing

Nucleic acid isolation, fragmentation, adapter ligation, and indexing were performed as previously described using the Qiagen AllPrep DNA/RNA Micro Kit and the SMARTer Stranded Total RNA-Seq Kit v2 (Takara) (46, 56). Sequencing was performed on an Illumina NextSeq 500 instrument, employing single-end sequencing with the NextSeq 500/550 V2 High Output reagent kit (1 x 75 cycles), and targeting a read depth of at least 10 x 10^6^ aligned reads per sample. Indexed samples were demultiplexed to fastq files with BCL2FASTQ v2.17.1.14, trimmed using Trimmomatic v0.38 (to remove end nucleotides with a phred score less than 30 while requiring a minimum length of 20 bp), and aligned to the mm10 (GRCm38) reference genome using TopHat v.2.1.0 (46). Counts data for uniquely mapped reads over exons were obtained using SeqMonk v1.45.4 and filtered to protein-coding genes and genes with at least 1 count per million in at least 2 samples. Differential gene expression analysis was performed with the edgeR v3.24.3 R/Bioconductor package using R v3.5.1 and RStudio v1.1.447 as stated in the text and individual figure legends and as described previously (54, 57). *K*-means clustering and heat maps were generated using the Morpheus web interface (https://software.broadinstitute.org/morpheus/). Gene Set Enrichment Analysis was performed using the GSEA v4.0.3 GSEAPreranked tool (58) with genes ordered by log2(fold-change) in average expression.

### Modified reduced representation bisulfite sequencing

Genomic DNA was subjected to mRRBS as previously described (46, 54, 59–62). Bisulfite conversion efficiency averaged 99.52% ± 0.052% (SD) as estimated by the measured frequency of unmethylated CpGs in λ-bacteriophage DNA (New England BioLabs, #N3013S) added at a 1:200 mass ratio to each sample. Processing and analysis were conducted as previously described using Trim Galore! v0.4.3, Bismark v0.16.3, the DSS v2.30.1 R/Bioconductor package, and quantified with the SeqMonk platform with the bisulphite feature methylation pipeline, all using the mm10 (GRCm38) reference genome. Well-observed CpGs were defined as in reference (62). Differentially methylated regions (DMR) were defined using the callDMR function within the DSS R/Bioconductor package using a *p*-value threshold of 0.05, a minimum length of 50 bp, a minimum significant CpG count of 2, merging DMR within 100 bp, and requiring 50% of the CpGs within a DMR to meet the significance threshold without a pre-defined methylation difference (delta). Functional enrichment analysis was performed using the GREAT tool (44) and the Mouse Genome Informatics Phenotype ontology (45) with the basal+extension association rule (constitutive 5 kb upstream and 1 kb downstream, up to 1000 kb max extension) against the whole genome as background. Metagene analysis was performed using SeqMonk’s quantitation trend plot function across Treg cell-specific super-enhancer elements (19) after liftover of coordinates from the mm9 to the mm10 reference genome (59).

### B16 melanoma model

100,000 B16-F10 cells (ATCC CRL-6475) were subcutaneously injected into the hair-trimmed flank of 8-12-week-old mice. All cells were determined to be free of *Mycoplasma* contamination before injection. Tumor progression was measured as previously described (46). A subset of tumors was resected post-mortem for photographic and flow cytometry analysis.

### Statistical analysis and reproducibility

*p*-values and FDR *q*-values resulting from two-tailed tests were calculated using statistical tests stated in the figure legends using GraphPad Prism v8.3.0 or R v3.5.1. Exact values are reported unless specified. Central tendency and error are displayed as mean ± standard deviation (SD) except as noted. Violin plots show median and quartiles. Numbers of biological and technical replicates are stated in the figures or accompanying legends. Histological and gel images are representative of at least two independent experiments. For next-generation sequencing experiments, indicated sample sizes were chosen to obtain a minimum of 10^6^ unique CpGs per biological replicate as modeled using the DSS statistical procedures (46, 54, 59–62). Computational analysis was performed using Genomics Nodes and Analytics Nodes on Quest, Northwestern University’s High-Performance Computing Cluster.

### Data availability

The raw and processed next-generation sequencing data sets have been uploaded to the GEO database (https://www.ncbi.nlm.nih.gov/geo/) under accession number GSE143974, which will be made public upon peer-reviewed publication.

### Code availability

Code used for RNA-seq processing is available from https://github.com/ebartom/NGSbartom.

Code used for mRRBS processing is available in the supplement to reference (62).

## Acknowledgements

LMN was supported by NIH award T32HL076139. MATA was supported by NIH award T32GM008152. KRA was supported by the David W. Cugell and Christina Enroth-Cugell Fellowship. BDS was supported by NIH awards K08HL128867 and U19AI135964 and the Francis Family Foundation’s Parker B. Francis Research Opportunity Award. The content is solely the responsibility of the authors and does not necessarily represent the official views of the funding sources. We wish to acknowledge the Northwestern University Flow Cytometry Core Facility supported by CA060553; the BD FACSAria SORP system was purchased with the support of S10OD011996. We also wish to acknowledge the Northwestern University RNA-Seq Center/Genomics Lab of the Pulmonary and Critical Care Medicine and Rheumatology Divisions. Histology services were provided by the Northwestern University Mouse Histology and Phenotyping Laboratory, which is supported by P30CA060553 awarded to the Robert H. Lurie Comprehensive Cancer Center. This research was supported in part through the computational resources and staff contributions provided by the Genomics Compute Cluster, which is jointly supported by the Feinberg School of Medicine, the Center for Genetic Medicine, and Feinberg’s Department of Biochemistry and Molecular Genetics, the Office of the Provost, the Office for Research, and Northwestern Information Technology. The Genomics Compute Cluster is part of Quest, Northwestern University’s high performance computing facility, with the purpose to advance research in genomics.

## Author Contributions

KAH, LMN, MATA, SEW, and BDS contributed to the conception, hypotheses delineation, and design of the study; KAH, LMN, MATA, KRA, S-YC, HA-V, YP, PC, MA, EMS, and BDS performed experiments/data acquisition and analysis; KAH, LMN, SEW, and BDS wrote the manuscript or provided substantial involvement in its revision.

---

**Supplemental Table 1.**
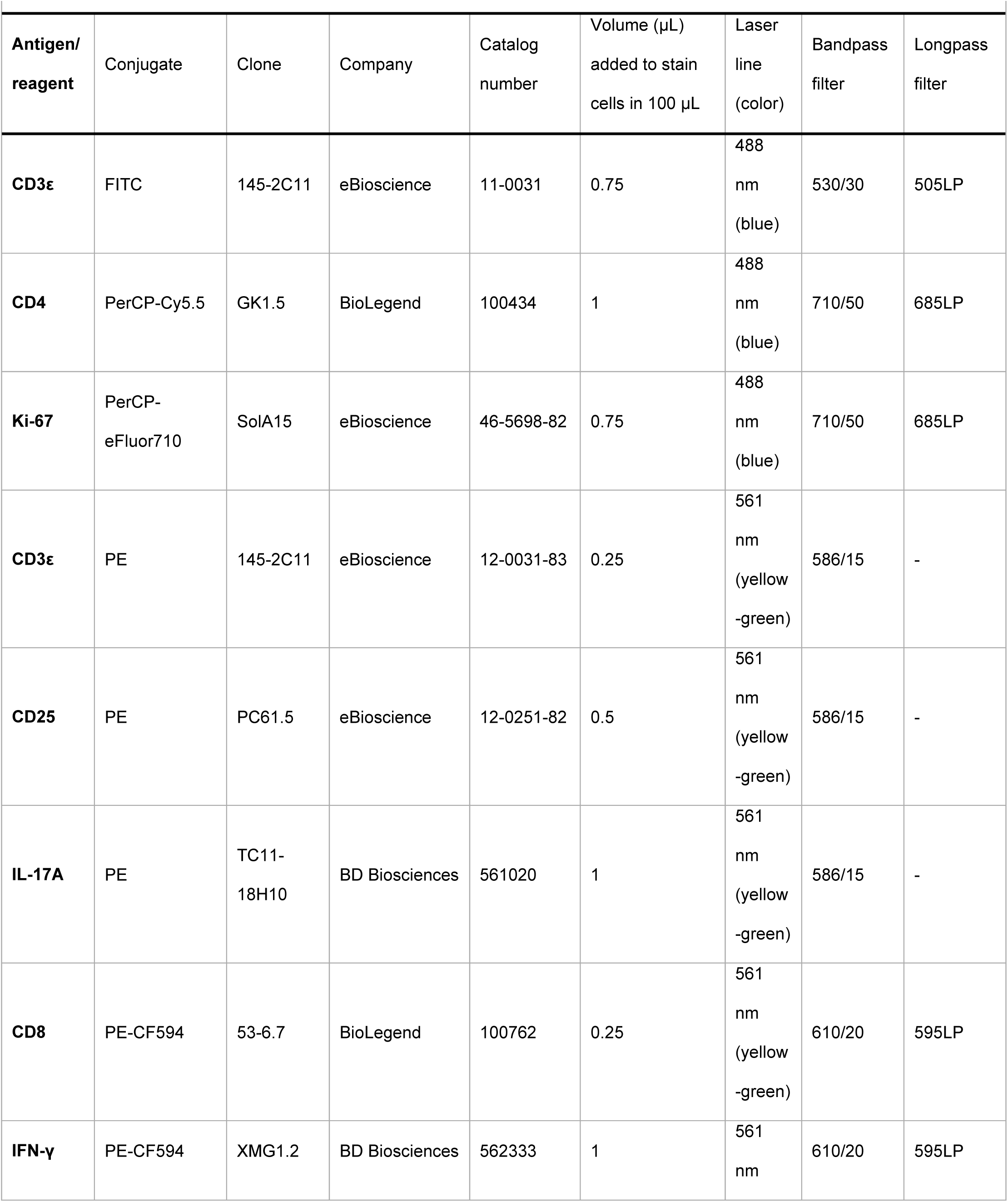
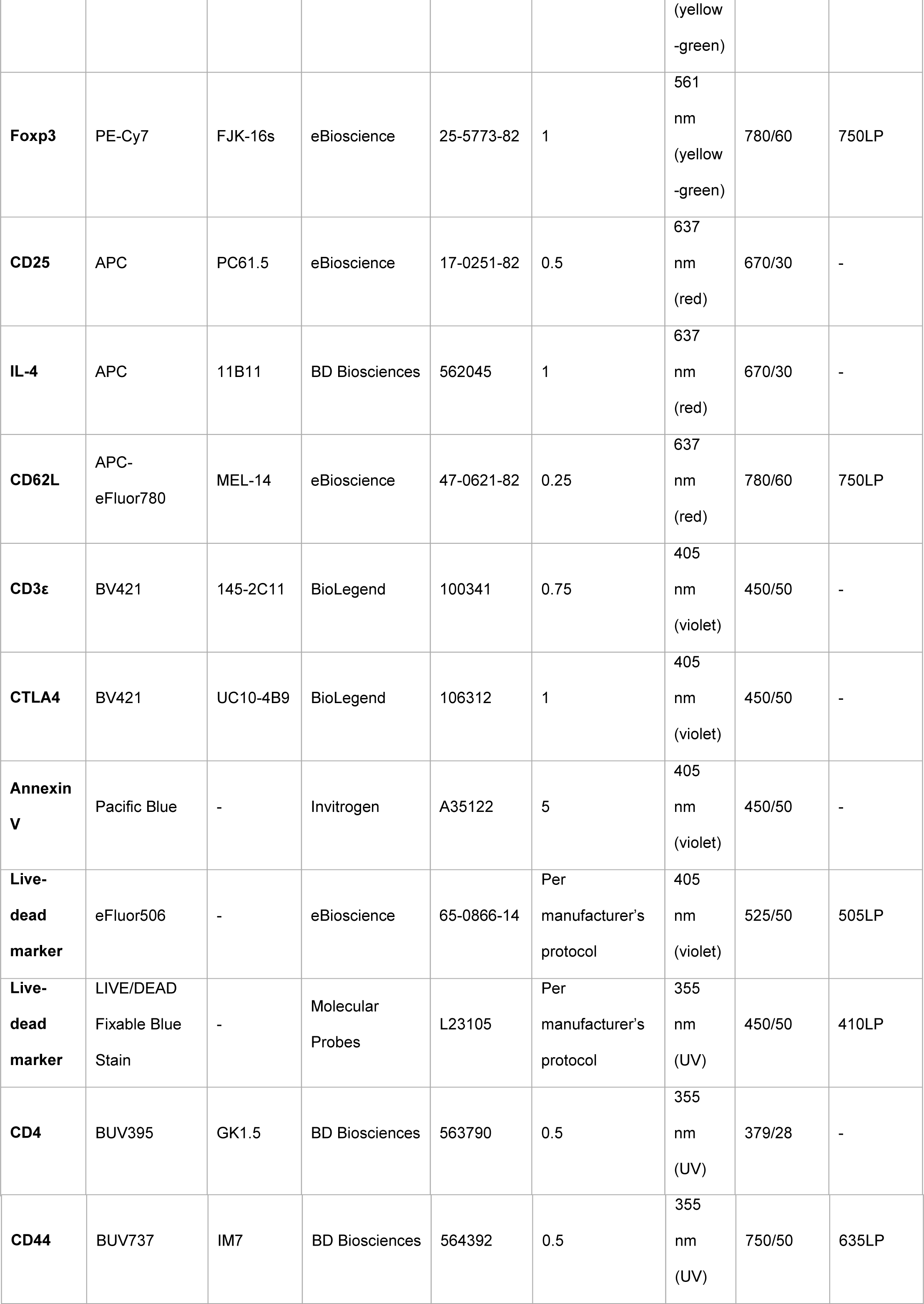
Reagents and cytometer setup. FITC, fluorescein isothiocyanate; PerCP, peridinin chlorophyll protein complex; PE, phycoerythrin; APC, allophycocyanin; BV, Brilliant Violet; BUV, Brilliant UltraViolet.

## References

1. Sakaguchi S, Yamaguchi T, Nomura T, Ono M. Regulatory T cells and immune tolerance. Cell. 2008;133(5):775–787.

2. Josefowicz SZ, Lu LF, Rudensky AY. Regulatory T cells: Mechanisms of differentiation and function. Annu Rev Immunol. 2012;30(1):531–564.

3. Singer BD, King LS, D’Alessio FR. Regulatory T cells as immunotherapy. Front Immunol. 2014;5:46.

4. Fontenot JD, Gavin MA, Rudensky AY. Foxp3 programs the development and function of CD4+CD25+ regulatory T cells. Nat Immunol. 2003;4:330–336.

5. Brunkow ME, et al. Disruption of a new forkhead/winged-helix protein, scurfin, results in the fatal lymphoproliferative disorder of the scurfy mouse. Nat Genet. 2001;27(1):68–73.

6. Wildin RS, et al. X-linked neonatal diabetes mellitus, enteropathy and endocrinopathy syndrome is the human equivalent of mouse scurfy. Nat Genet. 2001;27(1):18–20.

7. Valencia X, Yarboro C, Illei G, Lipsky PE. Deficient CD4+CD25high T regulatory cell function in patients with active systemic lupus erythematosus. J Immunol. 2007;178(4):2579–2588.

8. Valencia X, Lipsky PE. CD4+CD25+FoxP3+ regulatory T cells in autoimmune diseases. Nat Clin Pract Rheumatol. 2007;3(11):619–626.

9. Crispin JC, Martinez A, Alcocer-Varela J. Quantification of regulatory T cells in patients with systemic lupus erythematosus. J Autoimmun. 2003;21(3):273–276.

10. Miyara M, et al. Global natural regulatory T cell depletion in active systemic lupus erythematosus. J Immunol. 2005;175(12):8392–8400.

11. Morales-Nebreda L, Alakija O, Ferguson KT, Singer BD. Systemic lupus erythematosus-associated diffuse alveolar hemorrhage: A case report and review of the literature. Clin Pulm Med. 2018;25(5):166–169.

12. Antiga E, et al. Regulatory T cells in the skin lesions and blood of patients with systemic sclerosis and morphoea. Br J Dermatol. 2010;162(5):1056–1063.

13. Pardoll DM. The blockade of immune checkpoints in cancer immunotherapy. Nat Rev Cancer. 2012;12(4):252–264.

14. Peggs KS, Quezada SA, Chambers CA, Korman AJ, Allison JP. Blockade of CTLA-4 on both effector and regulatory T cell compartments contributes to the antitumor activity of anti-CTLA-4 antibodies. J Exp Med. 2009;206(8):1717–1725.

15. Simpson TR, et al. Fc-dependent depletion of tumor-infiltrating regulatory T cells co-defines the efficacy of anti-CTLA-4 therapy against melanoma. J Exp Med. 2013;210(9):1695–1710.

16. Jordan MS, et al. Thymic selection of CD4+CD25+ regulatory T cells induced by an agonist self-peptide. Nat Immunol. 2001;2(4):301–306.

17. Ohkura N, et al. T cell receptor stimulation-induced epigenetic changes and Foxp3 expression are independent and complementary events required for Treg cell development. Immunity. 2012;37(5):785–799.

18. Morales-Nebreda L, McLafferty FS, Singer BD. DNA methylation as a transcriptional regulator of the immune system. Transl Res. 2019;204:1–18.

19. Kitagawa Y, et al. Guidance of regulatory T cell development by Satb1-dependent super-enhancer establishment. Nat Immunol. 2017;18(2):173–183.

20. Singer BD, et al. Regulatory T cell DNA methyltransferase inhibition accelerates resolution of lung inflammation. Am J Respir Cell Mol Biol. 2015;52(5):641–652.

21. Lu CH, et al. DNA methyltransferase inhibitor promotes human CD4(+)CD25(h)FOXP3(+) regulatory T lymphocyte induction under suboptimal TCR stimulation. Front Immunol. 2016;7:488.

22. Chan MW, Chang CB, Tung CH, Sun J, Suen JL, Wu SF. Low-dose 5-aza-2’-deoxycytidine pretreatment inhibits experimental autoimmune encephalomyelitis by induction of regulatory T cells. Mol Med. 2014;20:248–256.

23. Kim H-P, Leonard WJ. Creb/atf-dependent T cell receptor-induced FoxP3 gene expression: A role for DNA methylation. J Exp Med. 2007;204:1543–1551.

24. Wang L, et al. Foxp3+ T-regulatory cells require DNA methyltransferase 1 expression to prevent development of lethal autoimmunity. Blood. 2013;121(18):3631–3639.

25. Samstein RM, et al. Foxp3 exploits a pre-existent enhancer landscape for regulatory T cell lineage specification. Cell. 2012;151(1):153–166.

26. Miyao T, et al. Plasticity of Foxp3(+) T cells reflects promiscuous Foxp3 expression in conventional T cells but not reprogramming of regulatory T cells. Immunity. 2012;36(2):262–275.

27. Zhou X, et al. Instability of the transcription factor Foxp3 leads to the generation of pathogenic memory T cells in vivo. Nat Immunol. 2009;10(9):1000–1007.

28. Ohkura N, Kitagawa Y, Sakaguchi S. Development and maintenance of regulatory T cells. Immunity. 2013;38(3):414–423.

29. Rubtsov YP, et al. Stability of the regulatory T cell lineage in vivo. Science. 2010;329(5999):1667–1671.

30. Bostick M, Kim JK, Esteve PO, Clark A, Pradhan S, Jacobsen SE. UHRF1 plays a role in maintaining DNA methylation in mammalian cells. Science. 2007;317(5845):1760–1764.

31. Sharif J, et al. The SRA protein Np95 mediates epigenetic inheritance by recruiting Dnmt1 to methylated DNA. Nature. 2007;450(7171):908–912.

32. Nishiyama A, et al. Uhrf1-dependent h3k23 ubiquitylation couples maintenance DNA methylation and replication. Nature. 2013;502(7470):249–253.

33. Kong X, et al. Defining UHRF1 domains that support maintenance of human colon cancer DNA methylation and oncogenic properties. Cancer Cell. 2019;35(4):633–648 e637.

34. Maenohara S, et al. Role of UHRF1 in de novo DNA methylation in oocytes and maintenance methylation in preimplantation embryos. PLoS Genet. 2017;13(10):e1007042.

35. Zhang J, et al. S phase-dependent interaction with DNMT1 dictates the role of UHRF1 but not UHRF2 in DNA methylation maintenance. Cell Res. 2011;21:1723.

36. Li E, Bestor TH, Jaenisch R. Targeted mutation of the DNA methyltransferase gene results in embryonic lethality. Cell. 1992;69(6):915–926.

37. Obata Y, et al. The epigenetic regulator Uhrf1 facilitates the proliferation and maturation of colonic regulatory T cells. Nat Immunol. 2014;15(6):571–579.

38. Sun X, Cui Y, Feng H, Liu H, Liu X. TGF-beta signaling controls Foxp3 methylation and T reg cell differentiation by modulating Uhrf1 activity. J Exp Med. 2019;216(12):2819–2837.

39. Rubtsov YP, et al. Regulatory T cell-derived interleukin-10 limits inflammation at environmental interfaces. Immunity. 2008;28(4):546–558.

40. Hill JA, et al. Foxp3 transcription-factor-dependent and -independent regulation of the regulatory T cell transcriptional signature. Immunity. 2007;27(5):786–800.

41. May SL, et al. Nfatc2 and Tob1 have non-overlapping function in T cell negative regulation and tumorigenesis. PLoS One. 2014;9(6):e100629.

42. Yan Y, et al. TCR stimulation upregulates MS4a4B expression through induction of AP-1 transcription factor during T cell activation. Mol Immunol. 2012;52(2):71–78.

43. Bonacci B, et al. Requirements for growth and IL-10 expression of highly purified human T regulatory cells. J Clin Immunol. 2012;32(5):1118–1128.

44. McLean CY, et al. GREAT improves functional interpretation of cis-regulatory regions. Nat Biotechnol. 2010;28(5):495–501.

45. Blake JA, Bult CJ, Eppig JT, Kadin JA, Richardson JE, Mouse Genome Database Group. The mouse genome database genotypes::Phenotypes. Nucleic Acids Res. 2009;37(Database issue):D712–719.

46. Weinberg SE, et al. Mitochondrial complex III is essential for suppressive function of regulatory T cells. Nature. 2019;565(7740):495–499.

47. Lal G, et al. Epigenetic regulation of Foxp3 expression in regulatory T cells by DNA methylation. J Immunol. 2009;182(1):259–273.

48. Yang R, et al. Hydrogen sulfide promotes Tet1- and Tet2-mediated Foxp3 demethylation to drive regulatory T cell differentiation and maintain immune homeostasis. Immunity. 2015;43(2):251–263.

49. Polansky JK, et al. DNA methylation controls Foxp3 gene expression. Eur J Immunol. 2008;38(6):1654–1663.

50. Varanasi SK, Reddy PBJ, Bhela S, Jaggi U, Gimenez F, Rouse BT. Azacytidine treatment inhibits the progression of herpes stromal keratitis by enhancing regulatory T cell function. J Virol. 2017;91(7).

51. Wu CJ, Yang CY, Chen YH, Chen CM, Chen LC, Kuo ML. The DNA methylation inhibitor 5-azacytidine increases regulatory T cells and alleviates airway inflammation in ovalbumin-sensitized mice. Int Arch Allergy Immunol. 2013;160(4):356–364.

52. Zhang Y, et al. Two missense variants in UHRF1BP1 are independently associated with systemic lupus erythematosus in Hong Kong chinese. Genes Immun. 2011;12(3):231–234.

53. Singer BD, et al. Flow-cytometric method for simultaneous analysis of mouse lung epithelial, endothelial, and hematopoietic lineage cells. Am J Physiol Lung Cell Mol Physiol. 2016;310(9):L796–801.

54. McGrath-Morrow SA, et al. DNA methylation regulates the neonatal CD4(+) T-cell response to pneumonia in mice. J Biol Chem. 2018;293(30):11772–11783.

55. Tighe RM, et al. Improving the quality and reproducibility of flow cytometry in the lung. An official American Thoracic Society workshop report. Am J Respir Cell Mol Biol. 2019;61(2):150–161.

56. Misharin AV, et al. Monocyte-derived alveolar macrophages drive lung fibrosis and persist in the lung over the life span. J Exp Med. 2017;214(8):2387–2404.

57. McGrath-Morrow SA, et al. Inflammation and transcriptional responses of peripheral blood mononuclear cells in classic ataxia telangiectasia. PLoS One. 2018;13(12):e0209496.

58. Subramanian A, et al. Gene set enrichment analysis: A knowledge-based approach for interpreting genome-wide expression profiles. Proc Natl Acad Sci U S A. 2005;102(43):15545–15550.

59. Walter JM, Helmin KA, Abdala-Valencia H, Wunderink RG, Singer BD. Multidimensional assessment of alveolar T cells in critically ill patients. JCI Insight. 2018;3(17):e123287.

60. Wang L, et al. TET2 coactivates gene expression through demethylation of enhancers. Sci Adv. 2018;4(11):eaau6986.

61. Piunti A, et al. CATACOMB: An endogenous inducible gene that antagonizes H3K27 methylation activity of polycomb repressive complex 2 via an H3K27M-like mechanism. Sci Adv. 2019;5(7):eaax2887.

62. Singer BD. A practical guide to the measurement and analysis of DNA methylation. Am J Respir Cell Mol Biol. 2019;61(4):417–428.

